# A digital twin of bacterial metabolism during cheese production

**DOI:** 10.1101/2023.05.05.539417

**Authors:** Maxime Lecomte, Wenfan Cao, Julie Aubert, David James Sherman, Hélène Falentin, Clémence Frioux, Simon Labarthe

## Abstract

Cheese organoleptic properties result from complex metabolic processes occurring in microbial communities. A deeper understanding of such mechanisms makes it possible to improve both industrial production processes and end-product quality through the design of microbial consortia. In this work, we caracterise the metabolism of a three-species community consisting of *Lactococcus lactis*, *Lactobacillus plantarum* and *Propionibacterium freudenreichii* during a seven-week cheese production process. Using genome-scale metabolic models and omics data integration, we modeled and calibrated individual dynamics using monoculture experiments, and coupled these models to capture the metabolism of the community. This digital twin accurately predicted the dynamics of the community, enlightening the contribution of each microbial species to organoleptic compound production. Further metabolic exploration raised additional possible interactions between the bacterial species. This work provides a methodological framework for the prediction of community-wide metabolism and highlights the added-value of dynamic metabolic modeling for the comprehension of fermented food processes.

## 1 Introduction

Fermentation of food and beverages by microbial consortia is one of the oldest food processing technologies, associated to health benefits, and used today in both traditional and industrial contexts (Tamang et al., 2016b,a). The transformation of raw food material by micro-organisms is the result of metabolic processes and interactions within the members of the microbial community. The content of the latter can be controlled, such as in industrial starters (Somerville et al., 2021), for food safety and for the predictability of organoleptic properties (Galimberti et al., 2021). While functional properties of microbial strains associated to fermented food get better understood (Tamang et al., 2016a), the precise molecular mechanisms at stake are only beginning to be elucidated (Blasche et al., 2021). In particular, the understanding of the individual metabolism of consortia members is not sufficient to capture or predict the global behaviour of the community system, which is impacted by microbe-microbe interactions.

Systems biology approaches combined to omics data acquisition are a promising avenue to characterise the functional landscape and interactions occurring in microbial consortia of fermented food (Mannaa et al., 2021). An approach on empirical communities can consist in combining metabolite and species inventories using culture-independent methods, and cultures of isolates (Blasche et al., 2021). For controlled community starters, a comparison of mono- and co-cultures can inform on the impact of interactions on the dynamics and metabolic profiles of the community Özcan et al. (2020). The use of genome-scale metabolic models (GEMs) (Fang et al., 2020) seems particularly opportune as a means to associate the strains of the consortia to the functions harboured by the systems and ultimately to the organoleptic properties of the food (Özcan et al., 2020). Such models of microbial consortia can be useful for numerous applications in food biotechnology, from the assembly of strains in starter design, to the optimization of processes for desired organoleptic properties (Rau and Zeidan, 2018).

The nature of the food matrix, the complexity of the bacterial community, and the highly strain-specific secondary metabolic pathways involved in the organoleptic features make it difficult to built mechanistic models, or digital twins of the biological system. Therefore, working with small-scale, controlled systems of reduced complexity is particularly relevant (Rau and Zeidan, 2018). Yet even simple systems must be considered and modeled over time, as fermentation is an intrinsically dynamic process. In this work, we are interested in modeling the metabolism of cheese making, from milk to ripening using a bacterial community starter mimicking industrial cheese production, composed of two Lactic Acid Bacteria (LAB), *Lactococcus lactis* and *Lactiplantibacillus plantarum*, and one propionic acid bacterium, *Propionibacterium freudenreichii*. Previous work (Cao et al., 2021) described how a fine-tuning of cheese standardised process parameters could modulate the production of aroma compounds. Here, using omics data, we build and exploit metabolic networks and community models in order (i) to evaluate the metabolic contribution of each microbial species during cheese making, (ii) to simulate the dynamics of the microbial metabolisms and the effect of the community over the individual behaviours, and (iii) propose possible interactions between the bacterial species.

Our work identifies the metabolic pathways responsible for nutrient transformation in milk. We optimized our individual models of metabolism and were able to reproduce the dynamics of pure cultures. From these models, we constructed a dynamic model of the bacterial community that accurately predicts the production of aroma compounds in the cheese.

## 2 Results

### 2.1 Individual metabolic models identify specific metabolic pathways in a milk environment

The microbial community starter was composed of *Propionibacterium freudenreichii* CIRM-BIA122, *Lactiplantibacillus plantarum* CIRM-BIA465 and *Lactococcus lactis* biovar *diacetylactis*. For each of them, full-length total DNA was obtained. Assembling bacterial genomes from a combination of short and long reads produced accurate and complete assemblies revealing one circular chromosome and the presence of 7 plasmids for the *L. lactis* strain and 4 plasmids for the *L. plantarum* strain. The *P. freudenreichii* strain did not harbour any plasmids. Annotated genome can be retrieved at ENA under accession ENA: PRJEB42478. The number of protein encoding genes, ribosomic sequences, tRNA and GC percent and accession numbers are gathered in Table 1. Annotation identified a list of 8234 coding sequences including those from chromosomal (2460 for *P. freudenreichii*, 2417 for *L. lactis* and 3016 for *L. plantarum*) and plasmid origins (139 for *L. lactis* and 82 for *L. plantarum*)

**Table 1:** Genomic overview of *P. freudenreichii*, *L. lactis* and *L. plantarum* after assembly.

|  |  |  |  |
| --- | --- | --- | --- |
| Bioproject Number | PRJEB42478 |  |  |
| Genome name | *Propionibacterium freudenreichii* | *Lactiplantibacillus plantarum* | *Lactococcus lactis* |
| Strain name | CIRM-BIA122 | CIRM-BIA465 | *lactis* biovar *diacetylactis*<br>CIRM-BIA1206 |
| Taxon id | 1744 | 1590 | 1358 |
| Accession number | ERS5564379 | ERS5564378 | ERS5564377 |
| number of plasmids | ∅ | 4 | 7 |
| size of plasmid (bases) | ∅ | p1: 40748, p2: 30463, p3: 9152, p4: 2012 | p1: 40748, p2: 30463, p3: 9152, p4: 2012, p5: 30463, p6: 9152, p7: 2012 |
| Size of the circular genome (bases) | 2622405 | 3121980 | 2365039 |
| number of rRNA | 6 | 16 | 19 |
| number of tRNA | 45 | 67 | 62 |
| percentage of GC (genome, min & max in plasmids) | 67 | 44 (36-41) | 35 (30-36) |
| number of CDS | 2472 | 3112 | 2563 |
| number of genes in each replicon | chromosome: 2557 | p1: 41, p2: 27, p3: 13, p4: 2, chromosome: 3157 | p1: 69, p2: 43, p3: 14, p4: 9, p5: 5, p6: 4, p7: 3, chromosome: 2542 |

Genome-scale metabolic networks were reconstructed for each individual strain starting from their respective annotated genomic sequences. After an automatic first step of reconstruction, we performed manual refinements to the central carbon metabolism based on literature and biological knowledge, in order to account for each strain’s specificity. The resulting metabolic networks satisfy the quality expectations in the field: a low fraction of universally-blocked reactions (*<* 3%), and a low fraction of reactions without gene-proteins-reactions rules (*<* 30%) (Lieven et al., 2020). Table 2 describes the characteristics of the three models. We observe that *L. lactis* has a smaller number of reactions than the other two strains, a qualitative difference that is also observed for other strains of the same species in other databases (Noronha et al., 2018).

**Table 2:** Characteristics of the three genome-scale metabolic network reconstructions and their associated FBA models in milk medium at the inoculation point.

|  | *P. freudenreichii* | *L. plantarum* | *L. lactis* |
| --- | --- | --- | --- |
| Number of genes | 1473 | 1433 | 1272 |
| Number of metabolites | 1284 | 1045 | 939 |
| Number of reactions | 1790 | 1523 | 1337 |
| Percentage of reactions associated to genes | 76.6 | 79.0 | 82.1 |
| Growth rate in mmol.gDW<sup>-1</sup>.hr<sup>-1</sup> | 0.187 | 0.645 | 2.172 |

We built genome-scale metabolic models (GEMs) from the three metabolic reconstructions using Flux Balance Analysis (FBA) (Palsson and Varma, 1994; Orth et al., 2010), ensuring the production of biomass nutrients on a nutritional environment reflecting milk composition (see Materials and Methods). We obtained growth rates ranging from 0.187 mmol.gDW*^−^*^1^.hr*^−^*^1^ to 2.17 mmol.gDW*^−^*^1^.hr*^−^*^1^, the former obtained for *P. freudenreichii* and the highest value obtained for *L. lactis*. Particular attention was given to the role of oxygen in the three metabolisms. Since bacterial growth on cheese corresponds to microaerobic conditions (Miyoshi et al., 2003), we altered the upper bounds of the oxygen transport reactions to limit their cytosolic import.

The central carbon metabolism produces NADH and NADPH reducing equivalents, ATP and precursors needed for the biosynthesis of molecules essential for the growth and the metabolic activities of bacterial cells e.g. amino acids, purine, pyrimidine, glycerol-3-phosphate, fatty acids, N-acetyl-glucosamine, vitamins. The three studied species shared almost the same glycolysis and pentose phosphate pathways. Carbon sources, in this case lactose, citric and lactic acids from milk, are converted into pyruvic acid, which is then metabolized *via* specific pathways. The specificity of each strain’s central carbon metabolism is illustrated in Figure 1. We describe below the characteristics of each strain together with the main refinements performed to their GEMs, paying specific attention on secondary pathways essential for organoleptic compound production.

**Figure 1:**
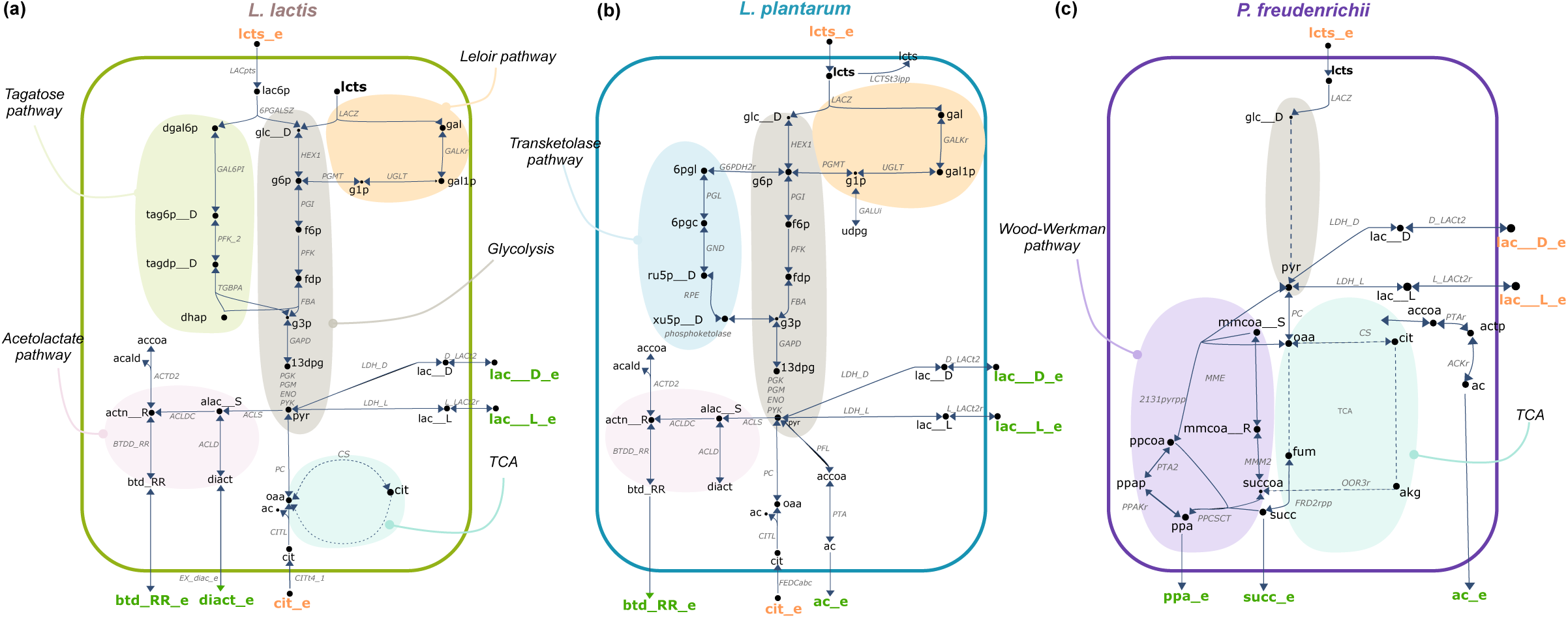
Metabolic maps of specific and shared pathways for *L. lactis* (a), *L. plantarum* (b) and *P. freudenreichii* (c). The glycolysis pathway (grey) is shared by the three species, while Leloir (salmon), acetolactate (pink) and citrate degradation pathways are shared by the two lactic acid bacteria. The TCA cycle (light blue) is shared by *P. freudenreichii* and *L. lactis*. Tagatose (green), transketolase (blue) and Wood-Werkman (purple) pathways are specific to *L. lactis*, *L. plantarum* and *P. freudenreichii* respectively. Orange and green metabolites are respectively inputs and outputs of the metabolic models. *Compound Abbreviations :* 13dpg: 3-Phospho-D-glyceroyl phosphate; 6pgc: 6-Phospho-D-gluconate; 6pgl: 6-phospho-D-glucono-1,5-lactone; ac: acetate; acald: acetaldehyde; accoa: acetyl-coa; actn R: acetoine; akg: 2-Oxoglutarate; alac S: (S)-2-Acetolactate; btd_RR: butanediol; cit: citrate; dgal6p: d-galactose-6-phosphate; dhap: Dihydroxyacetone phosphate; diact: diacetyl; f6p: fructose-6-phosphate; fdp: D-Fructose 1,6-bisphosphate; fum: fumarate; g1p: glucose-1-phosphate; g3p: Glyceraldehyde 3-phosphate; g6p: glucose-6-phosphate; gal: galactose; gal1p: galactose-1-phosphate; glc D: glucose; lac D, lac L: lactate; lac6p: lactose-6-phosphate; lcts: lactose; mmcoa R: (R)-Methylmalonyl-CoA; mmcoa S: (S)-Methylmalonyl-CoA; oaa: Oxaloacetate; ppa: propionate.; ppap: Propanoyl phosphate; ppcoa: propionyl-coa; pyr: pyruvate; ru5p D: D-Ribulose 5-phosphate; succ: succinate; succoa: succinyl-coa; tag6p D: D-tagatose-6-phosphate; tagdp D: D-Tagatose 1,6-biphosphate; udpg: UDPglucose; xu5p D: D-Xylulose 5-phosphate. *Reaction Abbreviations :* 2131pyrpp : Methylmalonyl-CoA carboxyltransferase 5S subunit; 6PGALSZ : 6-phospho-beta-galactosidase; ACLD : Acetolactate decarboxylase; ACTD2 : Acetoin dehydrogenase; ACLDC : Acetolactate decarboxylase; ACLS : Acetolactate synthase; BTDD_RR : R R butanediol dehydrogenase; CITL : Citrate lyase; CITt4_1 : Citrate transport via sodium symport; CS : Citrate synthase; D_LACt2 : D lactate transport via proton symport; ENO : Enolase; FBA : Fructose-bisphosphate aldolase; FEDCabc : FEDCabc; FRD2rpp : Fumarate reductase / succinate dehydrogenase (irreversible) (periplasmic, membrane potential dissipating); G6PDH2r : Glucose 6-phosphate dehydrogenase; GAL6PI : Galactose-6-phosphate isomerase; GALkr : Galactokinase; GALUi: UTP-glucose-1-phosphate uridylyltransferase (irreversible); GAPD : Glyceraldehyde-3-phosphate dehydrogenase; GND : Phosphogluconate dehydrogenase; HEX1 : Hexokinase (D-glucose:ATP); L_LACt2r : L lactate reversible transport via proton symport; LACpts : Lactose transport via PEP:Pyr PTS; LACZ : B-galactosidase; LCTSt3ipp : Lactose transport via proton aniport (periplasm); LDH_D : D-lactate dehydrogenase; LDH_L : L-lactate dehydrogenase; MME : Methylmalonyl-CoA epimerase; MMM2 : Methylmalonyl-CoA mutase; OOR3r : 2-oxoglutarate synthase (rev); PC : Pyruvate carboxylase; PFK : Phosphofructokinase; PFK_2 : Phosphofructokinase; PFL : Pyruvate formate lyase; PGI : Glucose-6-phosphate isomerase; PGK : Phosphoglycerate kinase; PGL : 6-phosphogluconolactonase; PGM : Phosphoglycerate mutase; PGMT : Phosphoglucomutase; phosphoketolase : phosphoketolase; PPAKr : Propionate kinase; PPCSCT : Propanoyl-CoA: succinate CoA-transferase; PTA : Phosphotransacetylase; PTA2 : Phosphate acetyltransferase; PYK : Pyruvate kinase; RPE : Ribulose 5-phosphate 3-epimerase; TGBPA : Tagatose-bisphosphate aldolase; UGLT : UDPgluc1o0se–hexose-1-phosphate uridylyltransferase

#### 2.1.1 LAB metabolism

Both *L. lactis* and *L. plantarum* utilize lactose and citric acid, the two main carbon sources in milk (Widyastuti et al., 2014). Lactose is degraded by a beta-galactosidase into glucose and galactose. Glucose feeds glycolysis, while galactose and citric acid metabolisms (Palles et al., 1998) are strain-dependent. Galactose is converted into glucose-6-phosphate *via*Leloir pathways before entering in glycolysis. Citric acid is converted into pyruvate by the citrate lyase multimeric enzyme encoded by the *citDEF* operon. Both species lead to the acidification of milk through the production of lactic and acetic acid (Roissart, 1994). The production of butanediol and diacetyl, responsible for a buttery flavour, occurs in both models through the acetolactate pathway from the transformation of pyruvate. All of these pathways are present and functional in the model (Fig. 1 a, b).

##### L. lactis

The production of butanediol required curation of the GEM (Fig. 1 a). The acetoin-dehydrogenase flux was blocked (*ACTD2*) permitting the production of butanediol, and bounds of the acetolactate decarboxylase (*ACLDC*), acetolactate synthase (*ACLS*) were refined in order to permit the activation of the acetolactate pathway, in accordance with the literature (Carroll et al., 1999; Swindell et al., 1996; Makhlouf, 2006). Finally, the consumption of lactose was regulated by refining the flux bounds of *LACZ* reaction which enabled a lactate production flux for *L. lactis* (Fig. 1 a).

To validate this model, we considered an example of pathway activation during cheese production. *L. lactis* consumes lactose either through the tagatose pathway, encoded in the *lacABCD* plasmidic operon, or by the Leloir pathway, encoded in the *galKTE* chromosomal operon. Metatranscriptomic data shows that *L. lactis* expresses the genes of the tagatose pathway at the beginning of cheese production, when lactose concentration is high, and those of the Leloir pathway during ripening, when lactose concentration is low. This strongly suggests that lactose concentration determines which pathway is employed, and predicts that a higher availability of lactose would produce a higher lactose flux through the tagatose pathway. The FBA model of *L. lactis* indeed leads to a greater flow of lactose through the tagatose pathway at the inoculation, and therefore meets the expectations (Fig. 2 a).

**Figure 2:**
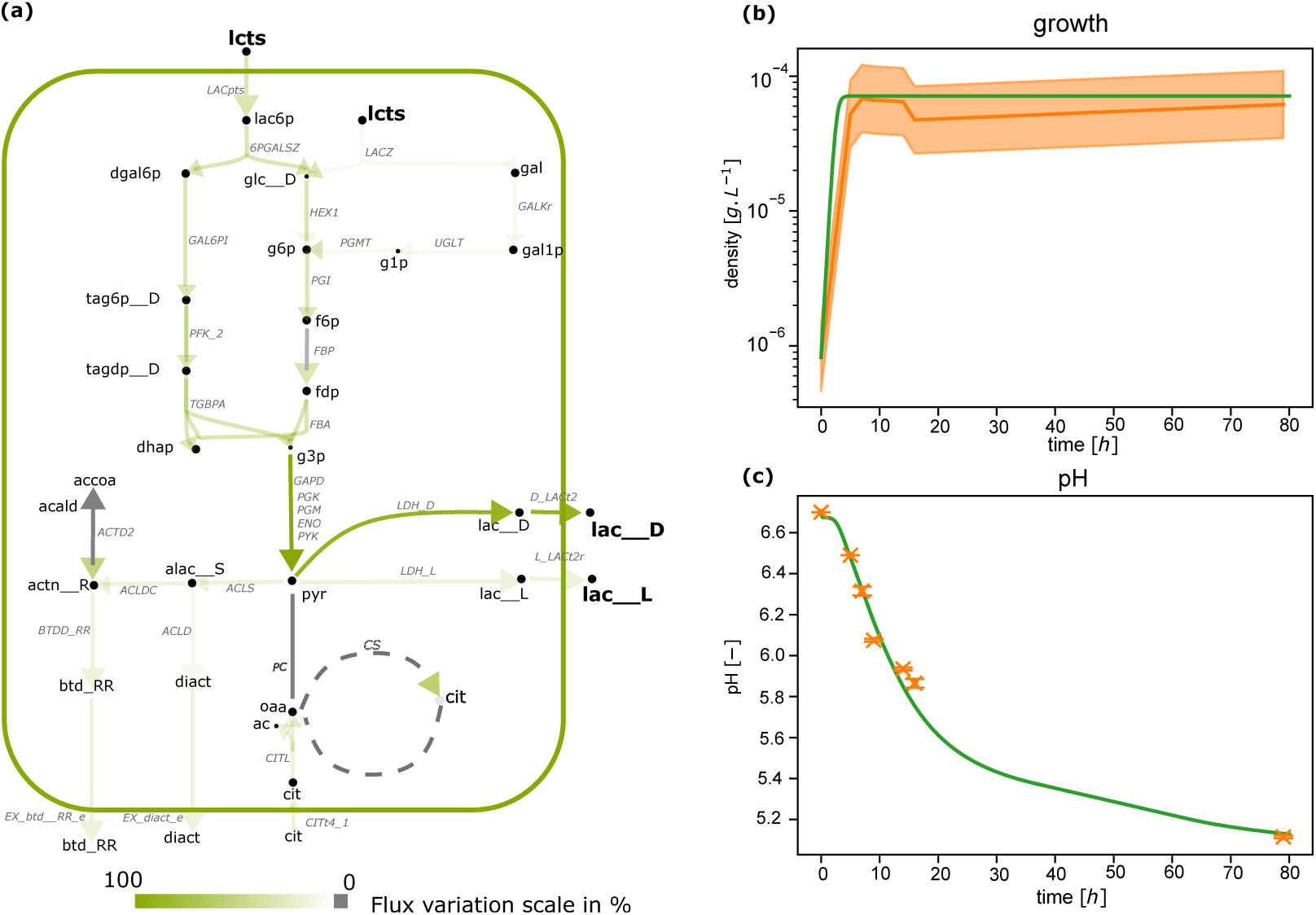
*L. lactis* GEM fitted on pure culture data. (a) FBA optimization with optimal parameters applied to the central carbon metabolism at the inoculation step. The color scale represents the reaction flux values predicted by the FBA and normalized by the highest flux value of the illustrated pathways. (b) Dynamics of *L. lactis* population in pure culture after parameter inference in the model (green line) and in the experimental data (orange line, with *±*1*/*4 log, which is the admitted range of precision for plating numbering). (c) pH in pure-culture of *L. lactis* in the model (green line) and in the experimental data (orange crosses, with standard deviation). *Compound abbreviations:* ac: acetate; actn R: acetoine; alac S: (S)-2-Acetolactate; btd_RR: butanediol; cit: citrate; dgal6p: d-galactose-6-phosphate; dhap: Dihydroxyacetone phosphate; diact: diacetyl; f6p: fructose-6-phosphate; fdp: D-Fructose 1,6-bisphosphate; g1p: glucose-1-phosphate; g3p: Glyceraldehyde 3-phosphate; g6p: glucose-6-phosphate; gal: galactose; gal1p: galactose-1-phosphate; glc D: glucose; lac D, lac L: lactate; lac6p: lactose-6-phosphate; lcts: lactose; oaa: Oxaloacetate; pyr: pyruvate; tag6p D: D-tagatose-6-phosphate; tagdp D: D-Tagatose 1,6-biphosphate. *Reaction abbreviations:* 6PGALSZ: 6-phospho-beta-galactosidase; ACLD: Acetolactate decarboxylase; ACTD2: Acetoin dehydrogenase; ACLDC: Aceto-lactate decarboxylase; ACLS: Acetolactate synthase; BTDD_RR: R R butanediol dehydrogenase; CITL: Citrate lyase; CITt4_1: Citrate transport via sodium symport; CS: Citrate synthase; D_LACt2: D lactate transport via proton symport; ENO: Enolase; FBA: Fructose-bisphosphate aldolase; GAL6PI: Galactose-6-phosphate isomerase; GALkr: Galactokinase; GAPD: Glyceraldehyde-3-phosphate dehydrogenase; HEX1: Hexokinase (D-glucose:ATP); L_LACt2r: L lactate reversible transport via proton symport; LACpts: Lac-tose transport via PEP:Pyr PTS; LACZ: B-galactosidase; LDH_D: D-lactate dehydrogenase; LDH_L: L-lactate dehydrogenase; PC: Pyruvate carboxylase; PFK: Phosphofructokinase; PFK_2: Phosphofruc-tokinase; PGI: Glucose-6-phosphate isomerase; PGK: Phosphoglycerate kinase; PGM: Phosphoglycerate mutase; PYK: Pyruvate kinase; TGBPA: Tagatose-bisphosphate aldolase; UGLT: UDPglucose–hexose-1-phosphate uridylyltransferase

##### L. plantarum

Glucose and galactose degradation lead to pyruvic acid which contributes to the release of lactic acid only in a homolactic metabolism or ethanol, acetic and lactic acids in a heterolactic metabolism. As *L. plantarum* is a heterolactic bacteria, it involves the specific transketolase enzyme (4.1.2.9), previously described in (Abedi and Hashemi, 2020); thus, acetic acid can be produced via the acetate kinase pathway. The production of butanediol required similar curation as for *L. lactis*. The conversion of pyruvate into acetic acid was ensured by blocking the pyruvate formate lyase (PFL) (Quatravaux et al., 2006). To reproduce the degradation of galactose and glucose through Leloir and glycolysis pathways by the *β*-galactosidase *LACZ*, the *LCTSt3ipp* reaction was knocked-out. To complete the galactose degradation, the UTP-glucose-1-phosphate uridylyltransferase (*GALUi*) was made reversible.

#### 2.1.2 P. freudenreichii metabolism

The CIRM-BIA122 strain of *P. freudenreichii* consumes either lactic acid, its main source of carbon (Thierry et al., 2011; Dank et al., 2021) or lactose (Loux et al., 2015). Lactic acid and lactose are catabolised through the glycolysis and the pentose-phosphate pathways, leading to pyruvic acid. One part of pyruvic acid fuels the tricarboxylic acid cycle (TCA) and the Wood-Werkman pathway (Fig. 1 c). Production of propionic acid, specific to the *Propionibacterium* genus, is ensured by routing succinic acid from TCA. The production of acetic acid and CO_2_ (Turgay et al., 2020) from pyruvate is permitted through the activity of two enzymes: the pyruvate dehydrogenase and the decarboxylative oxidation.

Ensuring the correct propionate-to-acetate ratio required further curation of the GEM. The bounds of *PPAKr, PTAr, 2131pyrpp* reactions were refined in order to drive propionate through the predominant production pathway, i.e. Wood-Werkman, and *POX2* reaction was blocked to regulate flux of acetate.

To validate the model, we computed the ratio of propionate/acetate and obtained a value of 2.19, consistent with biological observations of 1:2 (Crow, 1986; Dank et al., 2021; Cao et al., 2021; Turgay et al., 2020).

After refinement, FBA models of the three species predicted the production of several aroma compounds such as lactic, acetic, propionic acids, diacetyl, and butanediol (Smid and Kleerebezem, 2014; Cao et al., 2021) (Fig. 1). They furthermore accurately reproduce the metabolic specificities of each strain with respect to their primary metabolism, i.e. Wood-Werkman pathway for *P. freudenreichii*, tagatose pathway for *L. lactis* and transketolase pathway for *L. plantarum*.

### 2.2 Individual dynamic model optimization

Each microbial strain was cultivated in pure culture: monitoring of pH for LAB, and growth, for all strains, was performed in milk under microaerophilic conditions (Supp. Table S1). To mimic lactate availability in co-culture, the media was supplemented with lactate and a hydrolyzate of milk proteins simulating proteolysis for *P. freudenreichii* . Under these conditions, all strains reached a culturability above 9 log10 CFU/g and pH initially at 6.7 at inoculation time reached 5.7 and 5.1 in *L. plantarum* and *L. lactis* cultures respectively. The plateau of culturability corresponding to the entry into stationary phase was reached in 48 hours for *P. freudenreichii*, 14 hours for *L. plantarum* and 6 hours for *L. lactis*. We note the FBA models at the inoculation predicted the highest growth rate for *L. lactis*, and the lowest for *P. freudenreichii*, consistent with these experimental observations (Table 2). In a second experiment, metabolite dosage was performed for *P. freudenreichii* at the final time point, showing a large production of propionate (8g/L of milk), a strong consumption of lactate (9g/L of milk) and a lower production of acetate and succinate (respectively 3,07 g/L and 0.371g/L of milk) (Supp. Table S2).

We used the experimental cultures of the strains to calibrate dynamic models of their respective metabolism. Experimentally, we observed that the metabolism of LAB is still active at the steady-state. As lactic acid is produced from lactose, and lactose was not a limiting substrate, we therefore regulated the flux of lactic acid and acetate production by the flux of lactose consumption. In addition, following the approach of (Özcan et al., 2020), lactose consumption was further regulated by undissociated lactic acid (see Supplementary material §B.5). For *P. freudenreichii*, additional bounds on metabolite production were estimated on the acid dosage data described above in order to constraint production fluxes to physiological values (see Supplementary material §B.7). Furthermore, as carbon sources were not limited in the growth media, a logistic growth was assumed to model the plateau phase.

To calibrate the metabolic models, we estimated dynamic model parameters on these mono-culture data by solving the minimisation problem (8)-(9). To avoid over-fitting, a reduced number of free parameters was kept for each model. Namely, only two parameters were fitted for *L. lactis*, tuning pH-dependant regulation, three parameters for *L. plantarum* involved in pH-dependant regulation and logistic growth, and one parameter for *P. freudenreichii* tuning a logistic regulation (see Material and Methods §4.3 for details).

After optimization, the dynamic FBA (dFBA) simulations fitted with the experimental growth data (Fig. 2, 3 and 4, (b)). The pH curves (Fig. 2, 3, (c)) are also correctly rendered, showing a good agreement of the pH model, together with *P. freudenreichii* metabolite production (Fig. 4, (c)). An overall coefficient of determination of 0.99 was recovered (Supplementary data, Fig. S1, left panel).

**Figure 3:**
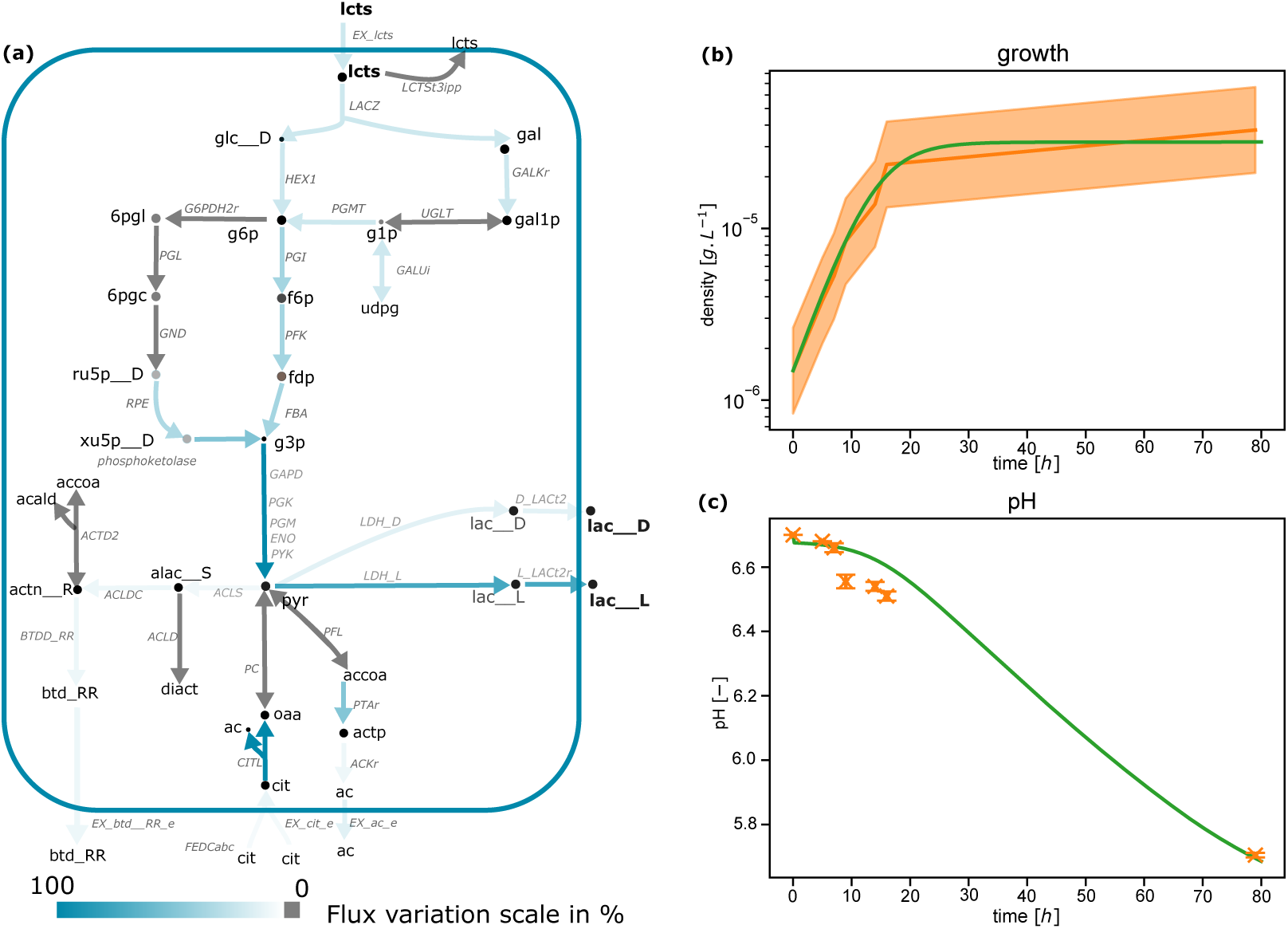
*L. plantarum* GEM fitted on pure culture data. (a) FBA optimization with optimal parameters applied to the central carbon metabolism at the inoculation step. The color scale represents the reaction flux values predicted by the FBA and normalized by the highest flux value of the illustrated pathways. (b) Dynamics of *L. plantarum* population in pure culture after parameter inference in the model (green line) and in the experimental data (orange line, with *±*1*/*4 log, which is the admitted range of precision for plating numbering). (c) pH in pure-culture of *L. plantarum* in the model (green line) and in the experimental data (orange crosses, with standard deviation). 6pgc: 6-Phospho-D-gluconate; 6pgl: 6-phospho-D-glucono-1,5-lactone; ac: acetate; acald: acetaldehyde; accoa: acetyl-coa; actn R: acetoine; alac S: (S)-2-Acetolactate; btd_RR: butanediol; cit: citrate; dhap: Dihydroxyacetone phosphate; f6p: fructose-6-phosphate; fdp: D-Fructose 1,6-bisphosphate; g1p: glucose-1-phosphate; g3p: Glyceraldehyde 3-phosphate; g6p: glucose-6-phosphate; gal: galactose; gal1p: galactose-1-phosphate; glc D: glucose; lac D, lac L: lactate; lcts: lactose; oaa: Oxaloacetate; pyr: pyruvate; ru5p D: D-Ribulose 5-phosphate; tag6p D: D-tagatose-6-phosphate; tagdp D: D-Tagatose 1,6-biphosphate; udpg: UDPglucose; xu5p D: D-Xylulose 5-phosphate. *Reaction abbreviations:* ACLD: Acetolactate decarboxylase; ACTD2: Acetoin dehydrogenase; ACLDC: Acetolactate decarboxylase; ACLS: Acetolactate synthase; BTDD_RR: R R butanediol dehydrogenase; D_LACt2: D lactate transport via proton symport; ENO: Enolase; FBA: Fructose-bisphosphate aldolase; FEDCabc: FEDCabc; G6PDH2r: Glucose 6-phosphate dehydrogenase; GALkr: Galactokinase; GALUi: UTP-glucose-1-phosphate uridylyl-transferase (irreversible); GAPD: Glyceraldehyde-3-phosphate dehydrogenase; GND: Phosphogluconate dehydrogenase; HEX1: Hexokinase (D-glucose:ATP); L_LACt2r: L lactate reversible transport via proton symport; LACZ: B-galactosidase; LCTSt3ipp: Lactose transport via proton aniport (periplasm); LDH_D: D-lactate dehydrogenase; LDH_L: L-lactate dehydrogenase; PC: Pyruvate carboxylase; PFL: Pyruvate formate lyase; PTA: Phosphotransacetylase; PGL: 6-phosphogluconolactonase; PFK: Phosphofructokinase; PGK: Phosphoglycerate kinase; PGI: Glucose-6-phosphate isomerase; PGM: Phosphoglycerate mutase; PGMT: Phosphoglucomutase; phosphoketolase: phosphoketolase; PYK: Pyruvate kinase; RPE: Ribulose 5-phosphate 3-epimerase; UGLT: UDPglucose–hexose-1-phosphate uridylyltransferase.

**Figure 4:**
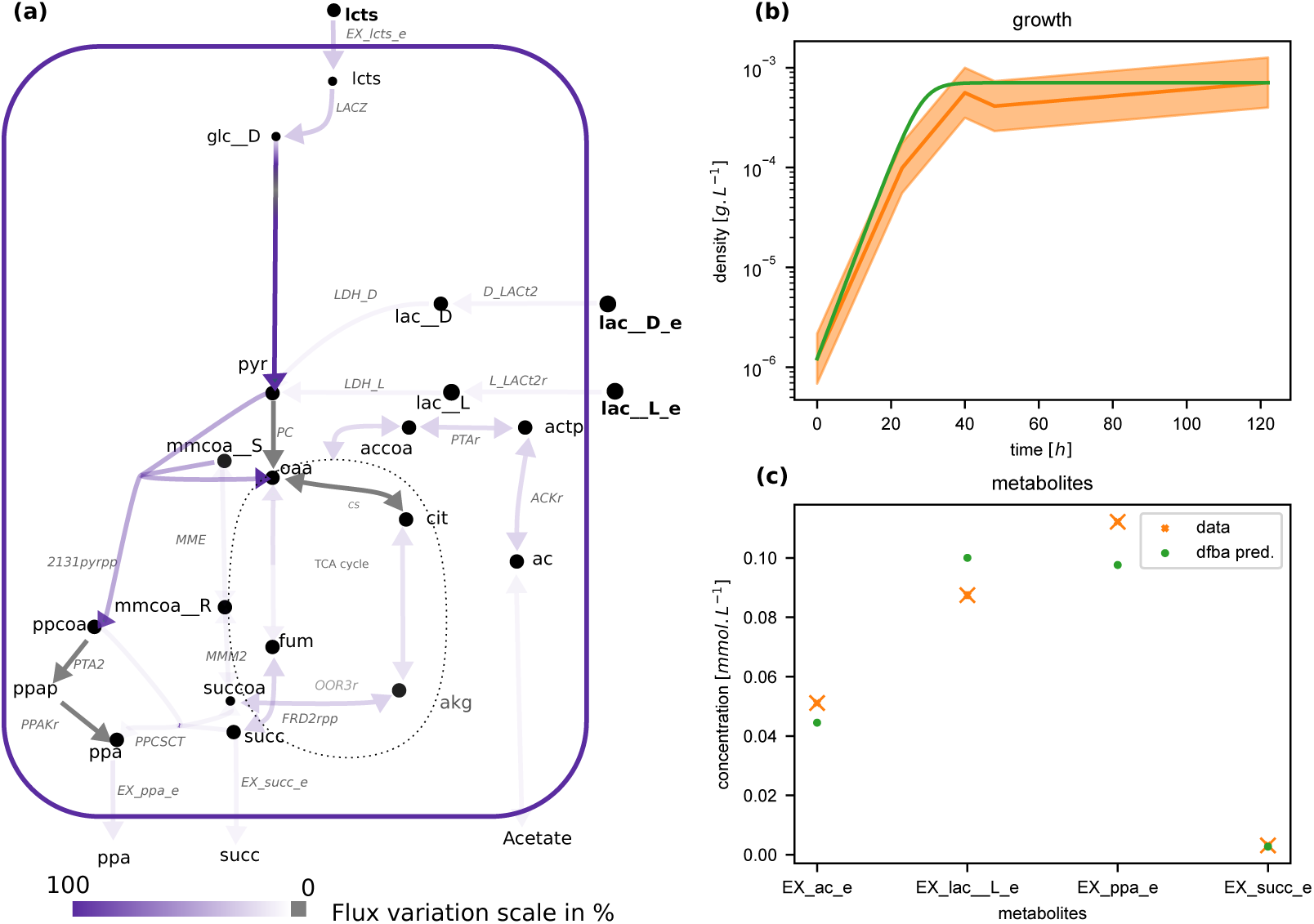
*P. freudenreichii* GEM fitted on pure culture data. (a) FBA optimization with optimal parameters applied to the central carbon metabolism at the inoculation step. The color scale represents the reaction flux values predicted by the FBA and normalized by the highest flux value of the illustrated pathways. (b) Dynamics of *P. freudenreichii* population size in pure culture after parameter inference in the model (green line) and in the experimental data (orange line, with *±*1*/*4 log, which is the admitted range of precision for plating numbering). (c) Lactate, acetate, propionate and succinate concentrations simulated (green points) and observed (orange crosses, with standard deviation) at the final time point of a second experiment with different lactate inoculation (see Material and method) *Compound abbreviations :* ac: acetate; akg: 2-Oxoglutarate; fum: fumarate; glc D: glucose; lac D, lac L: lactate; lcts: lactose; mmcoa R: (R)-Methylmalonyl-CoA; mmcoa S: (S)-Methylmalonyl-CoA; oaa: Oxaloacetate; ppa: propionate.; ppap: Propanoyl phosphate; ppcoa: propionyl-coa; pyr: pyruvate; succ: succinate; succoa: succinyl-coa.*Reaction abbreviations :* 2131pyrpp : Methylmalonyl-CoA carboxyltransferase 5S subunit; CS : Citrate synthase; D_LACt2 : D lactate transport via proton symport; FRD2rpp : Fumarate reductase / succinate dehydrogenase (irreversible) (periplasmic, membrane potential dissipating); L_LACt2r : L lactate reversible transport via proton symport; LACZ : B-galactosidase; LDH_D : D-lactate dehydrogenase; LDH_L : L-lactate dehydrogenase; MME : Methylmalonyl-CoA epimerase; MMM2 : Methylmalonyl-CoA mutase; OOR3r : 2-oxoglutarate synthase (rev); PC : Pyruvate carboxylase; PPAKr : Propionate kinase; PPCSCT : Propanoyl-CoA: succinate CoA-transferase; PTA2 : Phosphate acetyltransferase.

In order to validate that optimized FBA models correctly render known metabolism, fluxes of pathways of interest described in Figure 1 are represented in Figure 2, 3 and 4 (a) at inoculation, when no metabolite is limiting. The *L. lactis* model showed preferential fluxes of consumption of lactose *via* the tagatose pathway rather than Leloir pathway and production of L-and D-lactic acid *via* the glycolysis pathway, which explains the acidification of the milk and curd during cheese making (Fig. 2a). Production fluxes of diacetyl and butanediol occurred through an activation of acetolactate metabolism. An uptake flux of citrate by the *CITL* reaction encoded by the *citL* gene was observed.

The *L. plantarum* model presented a consumption flux of lactose and production fluxes of L-and D-lactic acid through the glycolysis pathway and acetic acid *via* the acetatekinase pathway, indicating that *L. plantarum* contributes to cheese acidification (Fig. 1b). The model shows different values of fluxes at different time points through the heterolactic (transketolase pathway) and homolactic (glycolysis pathway) metabolism. The activation of the transketolase pathway in a dairy context was, to our knowledge, observed for the first time. The consumption of citrate is strain dependant for *L. plantarum* (Palles et al., 1998) and was observed in our case through the *FEDCabc* transport reaction, leading citrate degradation to increase the pyruvate pool.

The *P. freudenreichii* model showed a higher consumption flux of lactose compared to D-lactic acid (Fig. 4a). TCA and Wood-Werkman cycles are activated simultaneously (Deborde and Boyaval, 2000): the former allows the regeneration of carbon source necessary for the activation of the Wood-Werkman cycle and thus, the production of propionate.

### 2.3 Follow-up of metabolites production in co-culture experiments by the community-wide model

Monitoring of growth and metabolic productions was performed during cheese production. *L. lactis*, *L. plantarum*, and *P. freudenreichii* reached respectively a culturability of 8.45 log_10_CFU/g, 8.47 log_10_CFU/g and 8.59 log_10_CFU/g in approximately 1200 hours. We built a community model by gathering the fitted individual models without further optimization, but the inclusion of the new plateau phase values in the logistic growth regulation in eq. 5, in order to reproduce *in silico* the metabolic behavior of the community (see eq. (3)-(4) and Material and methods §4.2). We note that, since no further fitting was performed, the co-culture experiments can be considered as a testing set involving unseen validation points.

The community model predicted the growth of each species quite accurately (Fig. 5(a)-(c), Supp. Table S4) despite a slight over-estimation of *P. freudenreichii* growth during the exponential phase. Metabolites dynamics were also correctly rendered. Lactose consumption is slower in the model compared to experiments, while citrate dynamics fits well with experimental data. Lactate production is slightly over-predicted during exponential phase, resulting in a slight over-acidification of the cheese during the molding and demolding steps. Acetate production is under-predicted during the early phases of cheese manufacture, but production rate during cheese maturation is correctly modeled. Curves for propionate, and to a lesser extent succinate, are accurately predicted by the model, suggesting that production of organoleptic compounds by *P. freudenreichii* is correctly captured. A global *R*^2^ of 0.98 is obtained for this validation step (Supplementary data, Figure S1, right panel). Furthermore, the metabolic pathways activated during mono-culture are also activated in co-culture (Supp. Fig. S3).

**Figure 5:**
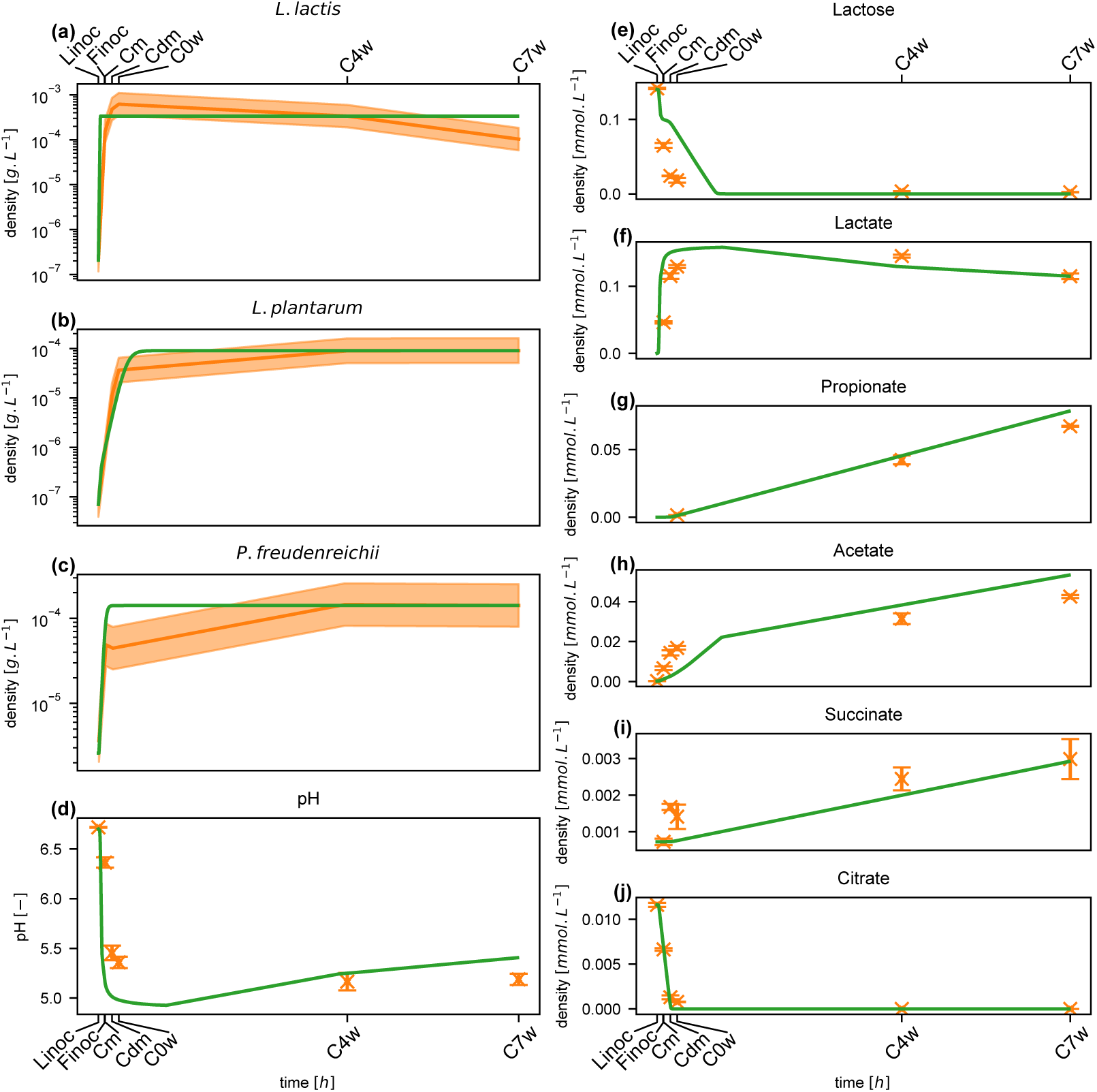
(a-c) Simulated growth of respectively *L. lactis*, *P. freudenreichii* and *L. plantarum* species computed by the community model (green lines) and experimentally-observed (orange lines, confidence interval of *±* 1/4 log). (d-j) Computational and experimental co-culture pH and metabolic profiles. Each data point is represented with its corresponding standard deviation (error bars). Abbreviations: Linoc, Lactic acid bacteria inoculation (t = 0); Finoc, *P. freudenreichii* inoculation (t = 18 hours); Cm, molding (t = 19.5 hours); Cdm, demolding (t = 40 hours); C0w, start of ripening (t = 60 hours); C4w, Fourth week ripening (t = 732 hours); C7w, seventh week ripening (t = 1236 hours).

### 2.4 Dynamical interactions within the community

Screening metabolite consumption and production during co-culture can reveal microbe-microbe interactions, such as cross-feeding or nutritional competition. We first surveyed the dynamics of metabolite exchange fluxes for each bacterium (Fig. 6, (a)) by computing, at each time step of the dynamics, the population-wide exchange flux *µ_i,j_b_i_* for metabolite *j* and bacterium *i* and the associated predicted metabolite concentrations. For citrate and lactose, *L. lactis* shows a strong uptake during its exponential phase, which rapidly vanishes after complete substrate depletion for citrate and activation of the pH-dependant regulation for lactose. Interestingly, *L. plantarum* fluxes are negligible for this substrate, suggesting that other metabolites support its growth, such as e.g. amino-acids. This may be due to the slower growth of *L. plantarum* during the first hours of cheese manufacture: citrate is depleted and acidification activates lactose consumption down-regulation when *L. plantarum* is still at low density. Lactose depletion is achieved by *P. freudenreichii* during the early phase of cheese ripening.

**Figure 6:**
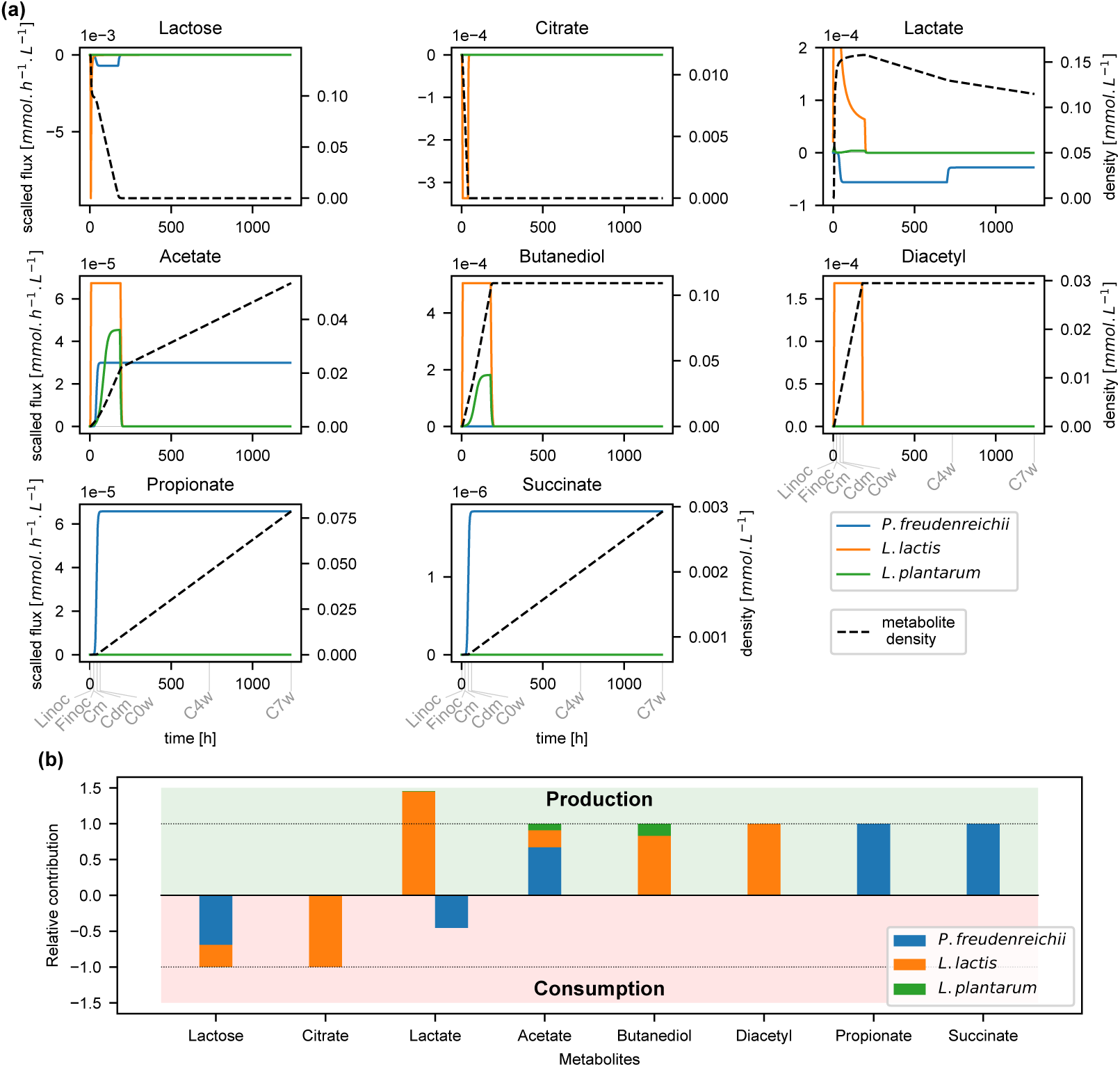
(a) **Consumption and production flux dynamics.** We represent for each bacterium the dynamics of the consumption and production fluxes of the followed-up metabolites (colored plain lines). We also represent the dynamics of the concentration of the corresponding metabolite in the cheese (dashed black line, right axis for scale). (b) **Bacterial relative contribution to the metabolite fate.** We represent the relative contribution of each bacteria to the final metabolite concentration. Namely, we c_f_omputed each microbial overall contribution by integrating in time its consumption or production flux (*^t^ µ_i,j_ b_i_* for metabolite *j* and bacteria *i*, see eq.4), and normalized the result by the sum over the three bacteria. The value 1 (or -1) then represents the net production (or consumption), i.e. the difference between the final and initial concentration of the metabolite (horizontal dashed lines). For each metabolite, the bacterial contribution is represented by a colored bar, with positive value for production, and negative value for consumption. Abbreviations: Linoc, Lactic acid bacteria inoculation (t = 0); Finoc, *P. freudenreichii* inoculation (t = 18 hours); Cm, molding (t = 19.5 hours); Cdm, demolding (t = 40 hours); C0w, start of ripening (t = 60 hours); C4w, Fourth week ripening (t = 732 hours); C7w, seventh week ripening (t = 1236 hours).

Lactate production reflects lactose consumption: lactate is strongly produced by *L. lactis* populations until *t* = 250*h* (ripening), while only a slight production by *L. plantarum* can be observed. Lactate is however consumed by *P. freudenreichii* during ripening, which contributes to the pH increase of the system observed in Figure 5 d. Acetate is produced by the three bacteria: *L. lactis* shows a strong acetate production until *t* = 250*h*, while *L. plantarum*’s contribution increases slowly until a plateau phase and drops at *t* = 250*h*. After reaching its plateau, *P. freudenreichii* keeps a constant rate of acetate production during cheese maturation. The same pattern is observed for butanediol production by the two LAB. Diacetyl is only produced by *L. lactis*, and its production stops after lactose depletion. Propionate and succinate are produced by *P. freudenreichii*, which keeps a constant release rate during ripening.

For each strain, we additionally computed the total net exchange fluxes of metabolites by integrating over time the FBA prediction 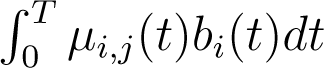. After normalization of this net exchange by the total exchange among the community (i.e. the sum of the individual net exchanges), we obtained the contribution of each strain to the metabolite fate (Fig. 6, (b)). These contributions confirm that succinate and propionate are solely produced by *P. freudenreichii*, while diacetyl (production) and citrate (consumption) are metabolized by *L. lactis*. *L. plantarum* contributes to butanediol production, although the main producer is predicted to be *L. lactis*, and to a smaller extent to acetate production, together with *L. lactis* and *P. freudenreichii* . Crossfeeding is observed for lactate between *L. lactis* (producer) and *P. freudenreichii* (consumer). Lactose consumption is shared between *L. lactis* and *P. freudenreichii* : interestingly, *P. freudenreichii* starts consuming lactose after *L. lactis* stops metabolising this substrate, indicating a time segregation of lactose use, and no direct competition for this substrate.

The above model demonstrates that capturing the complexity of metabolic processes in the bacterial community during cheese making requires consideration of the underlying dynamics occurring in the system. We wondered whether applying *a priori* computational approaches relying on genome-scale metabolic models could highlight additional putative interactions between the strains. To that end, we used two flux-based methods, namely SMETANA (Zelezniak et al., 2015) and MICOM Diener et al. (2020a), and one reasoning-based approach, Metage2Metabo (Belcour et al., 2020) in order to suggest metabolic complementarity among the consortium.

MICOM and SMETANA highlighted respectively 14 and 25 putatively exchanged metabolites (Supp. Table S5 and S6), while Metage2Metabo identified 11 metabolites that could not be produced by a species without interactions among the community (Supp. Table S7). A first observation is that a relatively high number of metabolites are common in the predictions of exchangeable compounds of SMETANA and MICOM: lactate, which was also predicted in the dynamic model, phenylalanine, serine, malate, succinate, xanthine, H_2_S, 2-ketoglutarate, glycollate and acetaldehyde. Other predicted exchanges include mostly additional amino-acids (isoleucine, proline, glycine, alanine). We used metatranscriptomic data in order to assess the validity of the most plausible interactions according to the SMETANA score (see Methods): H_2_S (from *L. lactis* and *L. plantarum* to *P. freudenreichii*), ribose (from *L. lactis* and plantarum to *P. freudenreichii*), glycerol (from *L. lactis* and *L. plantarum* to *P. freudenreichii*) and phenylanaline (from *L. plantarum* to *P. freudenreichii*). We verified the expression of genes associated to the production of the metabolite in the donor species, and to the consumption of the compound in the receiver species (see Methods and Supp. Fig. S2). Results suggest that interactions for H_2_S, ribose and glycerol are plausible at several steps of cheese making. Conversely, whereas gene expression data shows that phenylalanine consumption pathways are highly expressed in *P. freudenreichii*, their production by *L. plantarum* is not highly expressed suggesting that this interaction is less effective. Finally, the set of metabolites whose production is predicted by Metage2Metabo to require interactions among the community include various fatty-acids, as well as galactose-1-phosphate, benzyl-CoA, glyceraldehyde and xantosine, indicative of metabolic complementarity between the strains for the related metabolic pathways.

## 3 Discussion

In this work we provide a digital twin of the bacterial metabolism occurring in milk during cheese making. Using metabolic models of the three inoculated bacterial strains and multi-omics data, we were able to accurately reproduce metabolite production patterns and the dynamics of the bacterial populations. We relied on pure culture experiments to calibrate the individual metabolic models with few parameters for each, thereby limiting overfitting during community simulations. An originality of the model is its ability to predict the dynamics over the entire cheese making process (seven weeks). Our work supplies hypotheses on the underlying metabolic pathways at stake in the milk environment, as well as the contribution of each strain in the consumption of nutrients and the producibility of organoleptic compounds.

Knowledge from microbiology experts, scientific literature, and multiomics data were necessary to achieve our high-quality genome-scale metabolic models and subsequent dynamic simulations. The central carbon metabolism of LAB mainly produces lactic acid via glycolysis, tagatose and Leloir pathways from lactose degradation (Widyastuti et al., 2014; Van Rooijen et al., 1991; Kleerebezem et al., 2003); for the first time, citrate utilization and heterolactic fermentation were observed in the curated model of *L. plantarum* in milk. Ensuring the individual metabolic models can activate these pathways was therefore an important step of their validation, which enabled reproducing the enzymatic activities observed in (Quatravaux et al., 2006; Carroll et al., 1999). The propionic bacterium *P. freudenreichii* converts lactose, and preferentially lactate, into propionate according to (Loux et al., 2015; Thierry et al., 2011). Following (Borghei et al., 2021), we added the propionyl-CoA:succinate CoA transferase (2.8.3.-) enzyme for completing the Wood-Werkman cycle and thus enabling propionate production.

In (Özcan et al., 2020), individual tuning parameters of all media compounds are imposed in order to explain metabolomics at the species level, and to predict it at the community level. Compared to this work, we implemented a dynamic FBA based on (Mahadevan et al., 2002), that used strain specific parameters for predicting growth, pH and metabolic concentrations. To avoid over-fitting, we narrowed down the number of inferred parameters for each bacterium, keeping only two parameters tuning pH regulation for the LAB, and one parameter driving the growth for *L. plantarum* and *P. freudenreichii* . Additional calibration was conducted for *P. freudenreichii*, without any additional mathematical optimization, by computing upper bounds for metabolites from metabolomic data obtained in mono-culture.

The co-culture model gave insights into the community behaviour during cheese manufacture, and suggested temporality and contribution of each species to aroma coumpounds production. It showed that propionate and succinate production could only be attributed to *P. freudenreichii* in line with (Cao et al., 2021), and that diacetyl seems to be only produced by *L. lactis*. According to the community model, butanediol was produced during the moulding and at the beginning of the ripening by both *L. lactis* and *L. plantarum*, as observed in the respective mono-culture models. Regarding acetate production, the model infers an early production from moulding to ripening steps by *L. lactis* and *L. plantarum* followed by the major involvement of *P. freudenreichii* during ripening. Concerning carbon source utilization, while *L. plantarum* is completely equipped for lactose and citrate metabolisms, which are fully activated in mono-culture, it appears that these metabolic pathways are strongly down-regulated in co-culture. When cultivated with *L. lactis*, as *L. plantarum* growth is the slowest of both, it achieves its maximal metabolic efficiency after *L. lactis*’s exponential phase. As *L. lactis* mainly grows on lactose producing lactate, its exponential phase is associated with a strong acidification of the environment, which activates the pH-dependant repression of lactose metabolism in *L. plantarum*. *L. plantarum* may use amino-acids or other metabolites not followed-up by the dynamical model to support its growth. To further explore this hypothesis, we used community-scale metabolic model exploration tools to identify metabolic interactions outside the scope of the compounds we screened in the dynamical co-culture model, highlighting other putative metabolic interactions, of which a high proportion related to amino-acids.

In the co-culture model, *P. freudenreichii* keeps an active metabolism in the later phase of cheese manufacturing, during ripening. This metabolism is reflected by a lactate consumption all along the ripening phase, and a release of organoleptic compounds at a constant rate (Fig. 6). This behaviour is coherent with known metabolic capabilities of *P. freudenreichii* and with the metabolomics data obtained during cheese processing. In the model, *P. freudenreichii* also consumes the lactose remaining after *L. lactis* growth. Indeed, during its exponential phase, *L. lactis* mainly metabolizes lactose, strongly producing lactate and making the pH drop down. The pH reduction inhibits the pH-regulated lactose uptake in the two LABs, hence stopping lactose consumption by the *Lactobacilli*. Since lactose remains in the media and *P. freudenreichii* is the unique bacteria still capable to use it, it completes lactose depletion. In metatranscriptomics data, the LACZ gene is highly activated in *P. freudenreichii* during ripening, supporting lactose degradation by this strain after acidification (see Fig. S2).

This work highlights how the combination of expert knowledge, omic data and metabolic modeling can provide precise and accurate predictions on the mechanisms responsible for the dynamic behaviour of a microbial community. On the other hand, it also points out that the amount of data and the efforts necessary to create such high-quality models still remain an expensive price to pay despite the improvement of simulation approaches. Methodological development are yet to be proposed in order to automatize the calibration of models with data, and ensure both the correctness of the predictions and scalability to larger communities or communities of empirical composition.

## 4 Material and methods

### 4.1 Biological data

#### Community composition

Strains used in this project are described in (Cao et al., 2021). Briefly, the controlled bacterial community is composed of two lactic acid bacteria (LAB), *L. lactis* subsp. *lactis* biovar *diacetylactis* CIRM-BIA1206 (termed *L. lactis*), *L. plantarum* CIRM-BIA465 (termed *L. plantarum*), and one propionic bacterium, *P. freudenreichii* CIRM-BIA122 (termed *P. freudenreichii*) provided by the International Centre for Microbial Resources-Food Associated Bacteria (CIRM-BIA^1^).

#### Cheese manufacturing and sample collection

The standard cheese manufacture protocol described in (Cao et al., 2021) is used. Briefly, after collection of fresh milk, pasteurization (76*^◦^*C, 20 seconds), skimming, standardization (until 30 grams of fat and 36 mg of calcium per kg of milk, *L. lactis* and *L. plantarum* were inoculated at 5.7 and 5.2 log_10_ CFU*/*g (Colony Forming Unit). After prematuration of the inoculated milk (at 14 *^◦^*C for 18 h), *P. freudenreichii* was inoculated at 6.1 log_10_ CFU*/*g and stirred for 20 min. The milk was then poured in vats and warmed at 33*^◦^*C for 30 min. Commercial rennet was then added and coagulum was cut, stirred, washed, drained, prepressed and transferred into moulds for pressing. Demolding occurred on the third day of cheese making, and the cheeses were salted (immersion at 12*^◦^*C for 10h in saturated brine), dried overnight and vacuum-packed in plastic bags on the fourth day of cheese making for ripening during 7 weeks at 13 *^◦^*C.

After LAB bacteria inoculation (Linoc, t=0h) and *P. freudenreichii* inoculation (Finoc, t=18h), samples were collected at 5 different production stages for further analysis as described in (Cao et al., 2021): molding stage (Cm, t=19.5h), demolding stage (Cdm, t=40h), start of ripening (C0w, t=60h) and after 4 and 7 weeks of ripening (C4w, t=732h and C7w, t=1236h). Cheeses were replicated four times (biological repetitions realised different days with independant inocula and milks) and sampled each times for numeration and biochemistry and twice (biological repetitions) for RNA sequencing.

#### Single strain culture experiments

In order to calibrate metabolic models, a training dataset is built using single strain culture experiments. Single strain growth experiments of the three bacteria were performed in UHT full fat milk (Delisse) in tube under microaerobic conditions, i.e. without shaking in an incubator, at 30*^◦^*C, with two replicates. For *P. freudenreichii*, the milk was supplemented with lactate (20 g*/*L) and with casein peptone (10 g*/*L). A second growth experiment was performed for *P. freudenreichii* in order to measure metabolite levels (acetate, lactate, propionate and succinate) at the final time. *L. lactis* was inoculated at 1 *×* 10^6^and 4 *×* 10^6^ CFU*/*g, *L. plantarum* at 3 *×* 10^6^ and 6 *×* 10^6^ CFU*/*g and *P. freudenreichii* at 1.4 *×* 10^7^ and 9 *×* 10^6^ CFU*/*g. Pure culture growth data are available in Table S1.

#### Metatranscriptomic data

Ten grams of cheese were dispersed in 90g of a 2% sodium citrate solution during 1 min at max speed followed by 1 min at low speed in stomacher (Humeau, Treillière, France) at room temperature. Ten milliliters of the dispersion were centrifuged at 10 000 g at 4*^◦^*C during 5 min. Cell pellets were stored at *−*80*^◦^*C for RNA extraction. After defrost, the samples were centrifuged again for 5 min at 4*^◦^*C, 10 000 g to discard the supernatant. Cell pellets were suspended in 200 µL lysis buffer (50 mM Tris–HCl, 1 mM EDTA; pH 8.0) 20 mg*/*g lysozyme (MP Biomedicals, Illkirch, France) and 50 U*/*g mutanolysin (Sigma, Saint Quentin Fallavier, France) and incubated for 30 min at 24*^◦^*C. Suspensions were then transferred to 2 mL tubes containing 50 mg zirconium beads (diameter: 0.1 mm; BioSpec Products, Bartlesville) and 50 µL SDS (20 %). Samples were then shaken in a Precellys Evolution (Bertin, Montigny-le-Bretonneux, France) for two cycles of 40 s at 6500 rpm. RNA was then extracted from the cell lysates using Qiazol and RNeasy Mini kit (Qiagen) according to (Falentin et al., 2010) with two or three steps of subsequent DNase treatment (Dnase Rnase free, Ambion) according to the manufacturer’s instructions. RNA integrity (RIN) was evaluated using an Agilent 2100 bioanalyzer (Agilent Technologies, Santa Clara, CA). RNA concentrations were quantified using Qubit Fluorometric quantitation (Thermofisher, France). Lack of DNA contamination was also checked using Qbit and DNA quantitation protocol according to manufacturer instructions. RNA samples with a RIN value greater than 8, indicating a good RNA integrity, were kept for further analysis.

Metatranscriptomic data were obtained at five time steps of cheese manufacturing. Illumina libraries and sequencing were performed by GeneWiz (Leipzig, Germany). Reads are available at ENA (Cambridge) under accession number ENA: PRJEB42478. Sequenced reads were trimmed using Trim Galore and mapped against reference genomes with Bowtie 2 (Langmead et al., 2009). No mismatch was allowed and only reads that mapped to unique sequences were further analysed. Reads mapping to each coding sequence were counted using HTSeq-count (-stranded=reverse, -a 0, -t CDS, -i db_xref, -m union), (Putri et al., 2022).

Statistical analysis were conducted using R statistical software version 4.0.3 (R Core Team, 2021). RNA-Seq raw count data were normalized in two steps. First, a species-specific scaling factor was applied to eliminate composition biases between libraries. For each of the three species, they were calculated using all the genes of the considered species and two others rows for the other species calculated as the sum of the counts attributed to each of these two other species, using the method TMM (Trimmed Mean of M-Values) as implemented in the package edgeR version 3.32.1 (Robinson et al., 2009). A supplementary within-sample normalization step was performed to correct for gene length and enable comparison. Finally replicates were averaged.

#### Targeted metabolomics

Methods are available in (Cao et al., 2021). Briefly, sugars and organic acids in the samples were quantified using high performance liquid chromatography (HPLC). Volatile compounds were analysed using headspace (HS) trap extraction coupled to gas chromatography-mass spectrometry (GC-MS).

### 4.2 Metabolic modeling

#### Genome sequencing

Cell pellets (equivalent to 1e^10^ CFU) were obtained by centrifugation for 10 min at 5000 g from strain cultures in broth medium. DNA was extracted using DNeasy midi kits (Qiagen) according to the protocol described in (Falentin et al., 2010). Genomes were sequenced by Illumina NovaSeq 2 x 150 pb and PacBio Sequel (Eurofins Genomics, Constance, Germany). Genomes were assembled using Unicycler (Wick et al., 2017a,b) on the Galaxy Genotoul bioinformatics platform (Toulouse, France). Genome sequences were integrated in the MicroScope platform hosted at Genoscope (CEA, Evry, France) for automatic annotation according to (Vallenet et al., 2019). Annotated genomes are available at ENA (Cambridge) under accession number ENA: PRJEB54980.

#### Genome-scale metabolic model reconstruction

Individual bacterial genomes were functionally annotated using the MicroScope platform (Vallenet et al., 2019). Genome-scale metabolic network were then reconstructed from these annotated genomes with CarveMe(Machado et al., 2018) version 1.4.1. Each genome scale metabolic model was polished using ModelPolisher (Römer et al., 2016) which improved annotations by adding information related to BIGG identifiers from BiGG Models knowledgebase (King et al., 2015). The consistency of each model was checked with MEMOTE (Lieven et al., 2020) https://memote.io at the end of the polishing test. The medium composition included the minimal media found in CarveMe and compounds in milk https://ciqual.anses.fr/#/aliments/ 19024/lait-entier-pasteurise, with amino-acids were considered in excess (Supplementary File 1).

#### Flux balance analysis

A genome-scale metabolic model (GEM) was constructed for each strain using the metabolic network and the medium composition, and analysed using Flux Balance Analysis (FBA) (Orth et al., 2010) FBA computes an optimal flux distribution satisfying stoichiometric and biophysical feasibility providing maximal biomass, by solving the linear programming problem

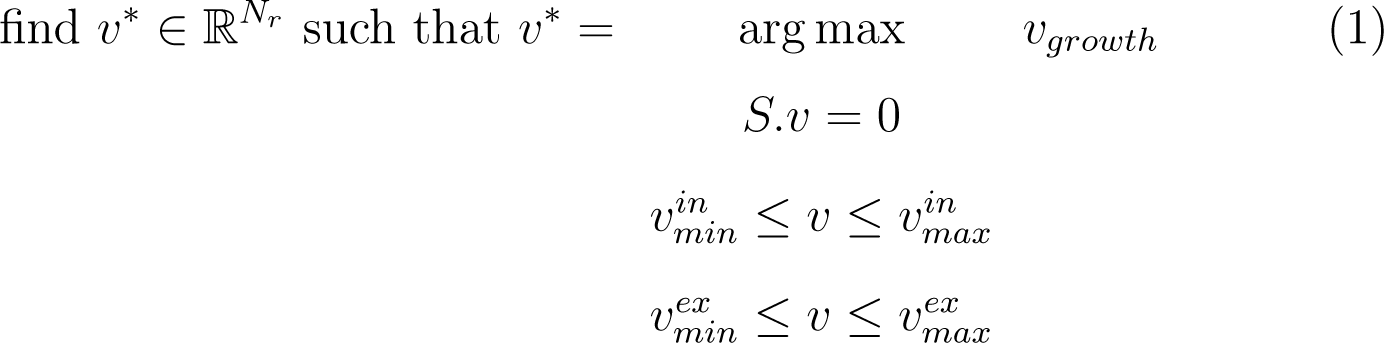

where *v_growth_* is the flux of the model biomass reaction, *S* is the stoichiometry matrix derived from the metabolic model, 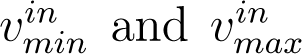 (resp. 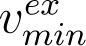 and 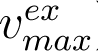) are minimal and maximal values for the intracellular (resp. exchange) fluxes defining flux bounds for the R*^Nr^* -dimensional flux vector *v*. The value of 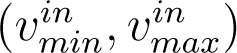 and 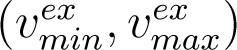 set in the GEMs are available in Supplementary Material.

Equation (1) then defines, for each bacterium *i*, the mapping *µ_i_*

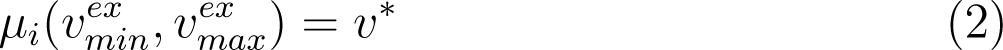

between the constraint vectors on exchange reactions defining the nutritional environment and the optimal flux *v^∗^* obtained in this nutritional context.

The FBA optimization problem (1) was solved using CobraPy (Ebrahim et al., 2013) version 0.17.1 using a defined lexicographic order (Gomez et al., 2014) to guarantee a unique solution.

#### GEM manual refinement

A manual refinement step was performed for all models to ensure organoleptic compound production and the activity of metabolic functions of interest with respect to literature knowledge. The resulting modifications can be found in Supplementary Material Table S3. As the three species are micro-aerobic, an uptake bound of oxygen is defined as the minimal value which permitted the production of aroma compounds.

#### Dynamic modeling

The dynamic behavior of each culture system was computed using dynamic flux balance analysis (dFBA) (Mahadevan et al., 2002) supplemented by additional population dynamics mechanisms.

Noting *b* = (*b_i_*) for *i ∈ B* = *{L. lactis, L. plantarum, P. freudenreichii}* the bacterial population densities, and *m* = (*m_j_*)_1_*_≤j≤Nm_* the density of the *N_m_* metabolic compounds that are experimentally screened, we set the system, for *i ∈ B* and 1 *≤ j ≤ N_m_*

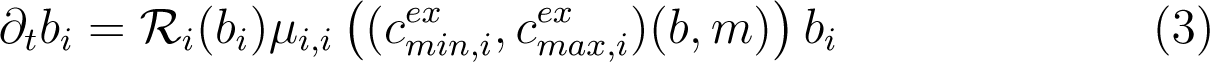

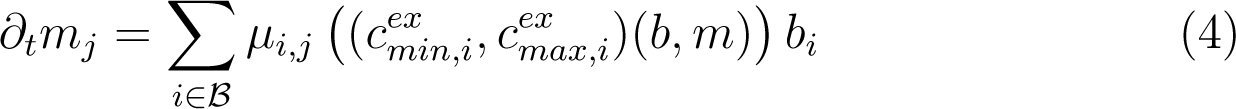

where the term *µ_i,j_* is the component corresponding to metabolite *j* (or biomass *i*) of *µ_i_*, computed from mapping 2 given the set of constraints 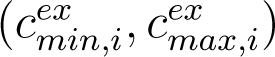 (*b, m*). These constraints depend dynamically on the state variables *b* and *m*.

The vector *R_i_* models a population-sensitive regulation process on the population growth in a phenomenological way with the term

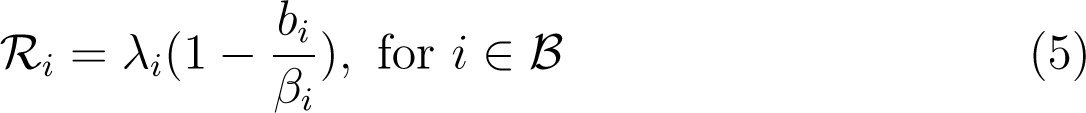

where *λ_i_* is a weight and *β_i_* is a population carrying capacity which is the plateau phase value of the corresponding bacteria in the experiments.

For a given substrate *j* and the metabolic model *i*, the usual bounds are

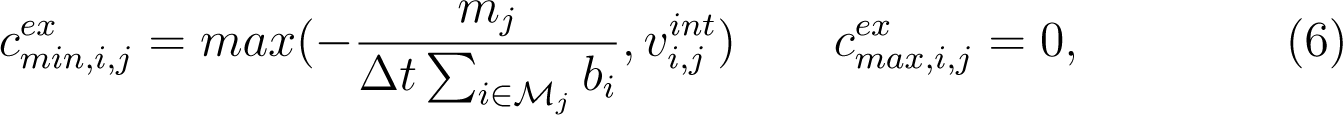

where *M_j_* is the subset of bacteria metabolizing *j* and Δ*t* is an import characteristic time. These equations reflect uniform sharing of resources among bacteria during the time window Δ*t*. When the substrate is in excess, the flux is limited by the intrinsic import limit 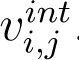.

Some substrates and products are additionally regulated. Following (Özcan et al., 2020), as the LAB bacteria metabolism of lactose is down-regulated by undissociated lactic acid with exponential decrease, for *j* = *lcts*_*e* and *i ∈ {L. lactis, plantarum}* we have

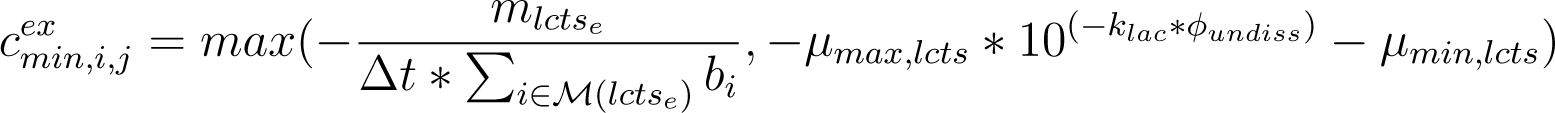

where *k_lac_* is an exponential decay, *µ_max,lcts_* and *µ_min,lcts_* are maximal and minimal values and *ϕ_undiss_* is a function computing the concentration of undissociated lactic acid from lactate. The function *ϕ_undiss_* is derived from Henderson–Hasselbalch equation as in (Özcan et al., 2020) and reads

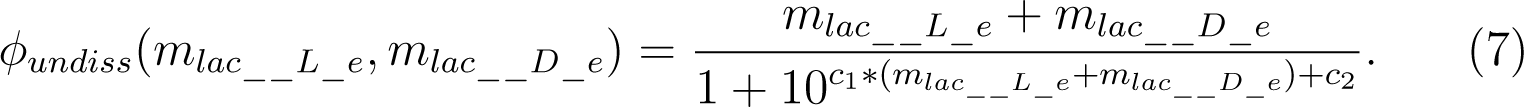

In this equation, the term *c*_1_ *∗* (*m_lac L_*___*_e_* + *m_lac D_*___*_e_*) + *c*_2_ is a linear approximation of (*pH − pKa*), so that parameters *c*_1_ and *c*_2_ are inferred by linear regression from data of the co-culture experiment. Lactate production is regulated in a similar way, while acetate production is regulated according to lactose availability. See Supplementary material sec. B.5 for a detailed description and justifications of the dynamical bounds set on the exchange reactions.

For *P. freudenreichii*, production upper bounds were evaluated by computing per capita production flux from the metabolite dosages and the growth data in the pure culture experiments, assuming a constant flux (see Supplementary material B.7).

The initial condition, the inoculum and nutritional environment composed of milk compounds, amino acids, and co-factors, was defined as specified in (Cao et al., 2021), after unit conversion (See supplementary material for a list of metabolites). A cell mass of 0.33 *×* 10*^−^*^12^ g*/*CFU was applied for biomass unit conversion.

#### Numerical implementation

The dynamical system is solved by a semi-implicit Euler scheme. At time step *n*, after computation of *F_j_* the total flux of compound *j* as defined by the set of GEMs,

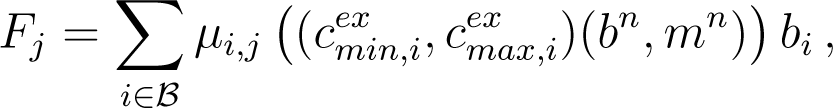

we computed

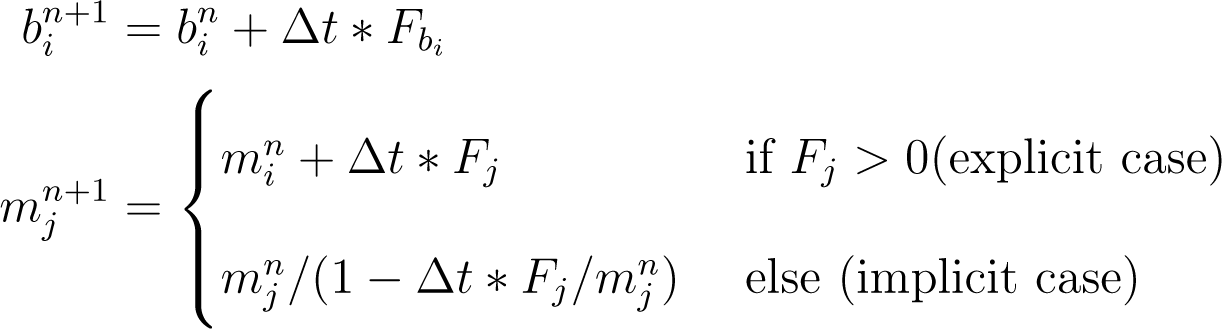

This scheme guaranties solution positivity at all times.

The dFBA problems (3)–(4) and parameter inference is solved using custom Python scripts and the scientific libraries numpy v.1.19.5(Harris et al., 2020), pandas v.0.25.3 (McKinney et al., 2010), and scipy v.1.5.3 (Wes McKinney, 2010). During dFBA, the FBA is solved using a defined lexicographic order (Gomez et al., 2014).

### 4.3 Model fitting

To fit the model parameters, we adapted equations (3)–(4) to mono-culture experiments. We then inferred the model parameter vector *θ* of the resulting dFBA model by maximising the log-likelihood of observed data in the monoculture. Namely, for *i ∈ {L. lactis, L. plantarum}*, we minimised the cost function

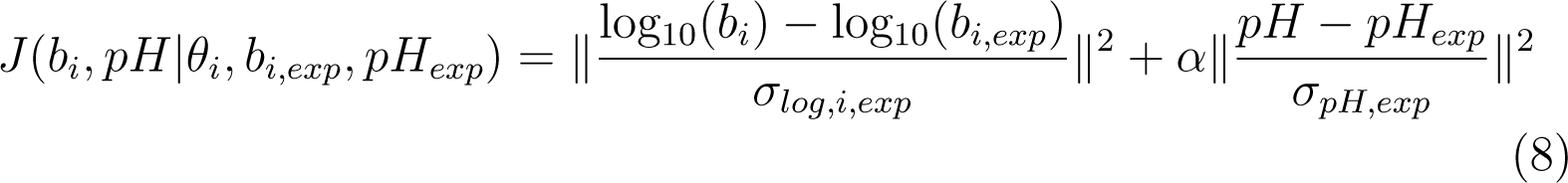

where *σ_i_* (resp. *σ_log,i_*) is the standard deviation (resp. the log-transformed standard deviation) of the corresponding data, and *θ_i_* = (*k_lac_, µ_max,lcts_*) if *i* = *L. lactis* and *θ_i_* = (*k_lac_, µ_max,lcts_, λ_i_*) if *i* = *L. plantarum*. For *P. freudenreichii*, the cost function is

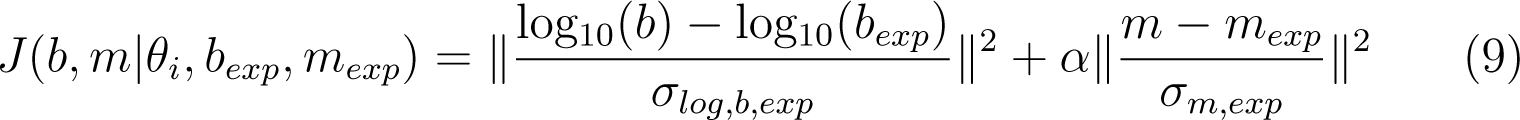

where *m_exp_* is the final dosage of acetate, lactate, propionate and succinate, and *θ_i_* = (*λ_i_*). We kept the number of fitted parameters small (2 parameters for *L. lactis*, 3 parameters for *L. plantarum* and 1 parameter for *P. freudenreichii*), to avoid over-fitting. Optimization results are presented in Table S8.

### 4.4 Inference of metabolic interactions

In order to predict metabolic exchanges likely to occur in the microbial consortium, MICOM v0.32.0 (Diener et al., 2020b) and SMETANA v1.2.0 (Zelezniak et al., 2015) were performed on the milk medium. Additionally, the prediction of metabolic complementarity between the three metabolic models grown in milk was performed with Metage2Metabo v.1.5.0 (Belcour et al., 2020). All metabolites predicted by the tools were classified into higher level categories: compounds identifiers were mapped to the Metacyc v.25.5 database (Caspi et al., 2018), and the ontology of the compounds was used. The list of predicted cross-feeding interactions is available in Supplementary Table S7.

A subset of putative bacterial interactions were selected among the ones predicted by SMETANA, according to their score (SMETANA score), for which a threshold was set at 0.5. The relevance of selected exchanged metabolites was assessed using metatranscriptomic data (RPKM and replicated means). For each metabolite, reactions producing (in the producer species) and consuming (in the consumer species) the metabolite were extracted from gene-protein-reactions associations. In order to evaluate the putative expression of these genes, the expression values were compared to the overall expression of genes in each species, separated into quartiles.

### 4.5 Visualization and statistical analyses

Plots were generated using matplotlib version 3.3.4 (Hunter, 2007). Metabolic maps for pathways of interest were generated with Escher (King et al., 2015) for Python v.3.6.9 (Van Rossum and Drake, 2009) ^2^, after running an FBA on each model. Fluxes of reactions of interest for each species were normalized by the maximal flux value of the targeted pathways in the corresponding species.

### 4.6 Code and data availability

Code for metabolic simulation and metabolic models is available in https://forgemia.inra.fr/tango/t Genomes of the three bacterial strains and their annotations are available in ENA under the accession number PRJEB54980. Metatranscriptomic data is available in ENA under the accession number PRJEB42478. Annotated genomes, RNA reads and RPKM are deposited in https://entrepot.recherche.data.gouv.fr/privateur 29dd-4d71-b25e-b775d9cc39dc. Biochemical dosage data and metabolomic data are available in (Cao et al., 2021).

## Acknowledgements

The authors acknowledge the Galaxy Genotoul bioinformatics platform (Toulouse, France), the MicroScope platform hosted at Genoscope (CEA, Evry, France), and the GenOuest bioinformatics core facility (https://www.genouest.org) for providing the computing infrastructure. CF, DJS and SL were supported by the French National Research Agency (ANR) France 2030 PEPR Agroécologie et Numérique MISTIC ANR-22-PEAE-0011. CF and SL were supported by the Inria Exploratory Action SLIMMEST. We are grateful to the INRAE STLO Dairy Platform (https://www6.rennes.inrae.fr/plateforme_lait_eng/) for providing help and support (doi: 10.15454/1.557240525786479E12), to the CNIEL for partial funding and fruitful discussions on the interpretation of the results (TANGO Project).

## 6 Authors contributions

Conceptualization: CF, HF, ML, SL; Data curation: JA, HF, ML; Formal analysis: CF, DJS, HF, ML,JA, WC, SL; Funding acquisition: HF; Investigation: CF, HF, ML, SL; Methodology: CF,HF, JA, ML, SL; Project administration: CF, DJS, HF, SL ; Resources: HF; Software: CF, ML, SL; Supervision: CF, DJS, HF, SL; Validation: HF, CF, ML, SL; Visualization: CF, ML, SL ; Roles/Writing -original draft: CF, HF, ML, SL; Writing -review & editing: all.

## 7 Supplementary information

The manuscript includes Supplementary Figures, Supplementary Material and a Supplementary File entitled *supplementary_file1.xlsx*.

## 8 Copyright

A CC-BY public copyright license has been applied by the authors to the present document and will be applied to all subsequent versions up to the Author Accepted Manuscript arising from this submission, in accordance with the grant’s open access conditions.

## 9 Bibliography

## A Supplementary figures

**Figure S1:**
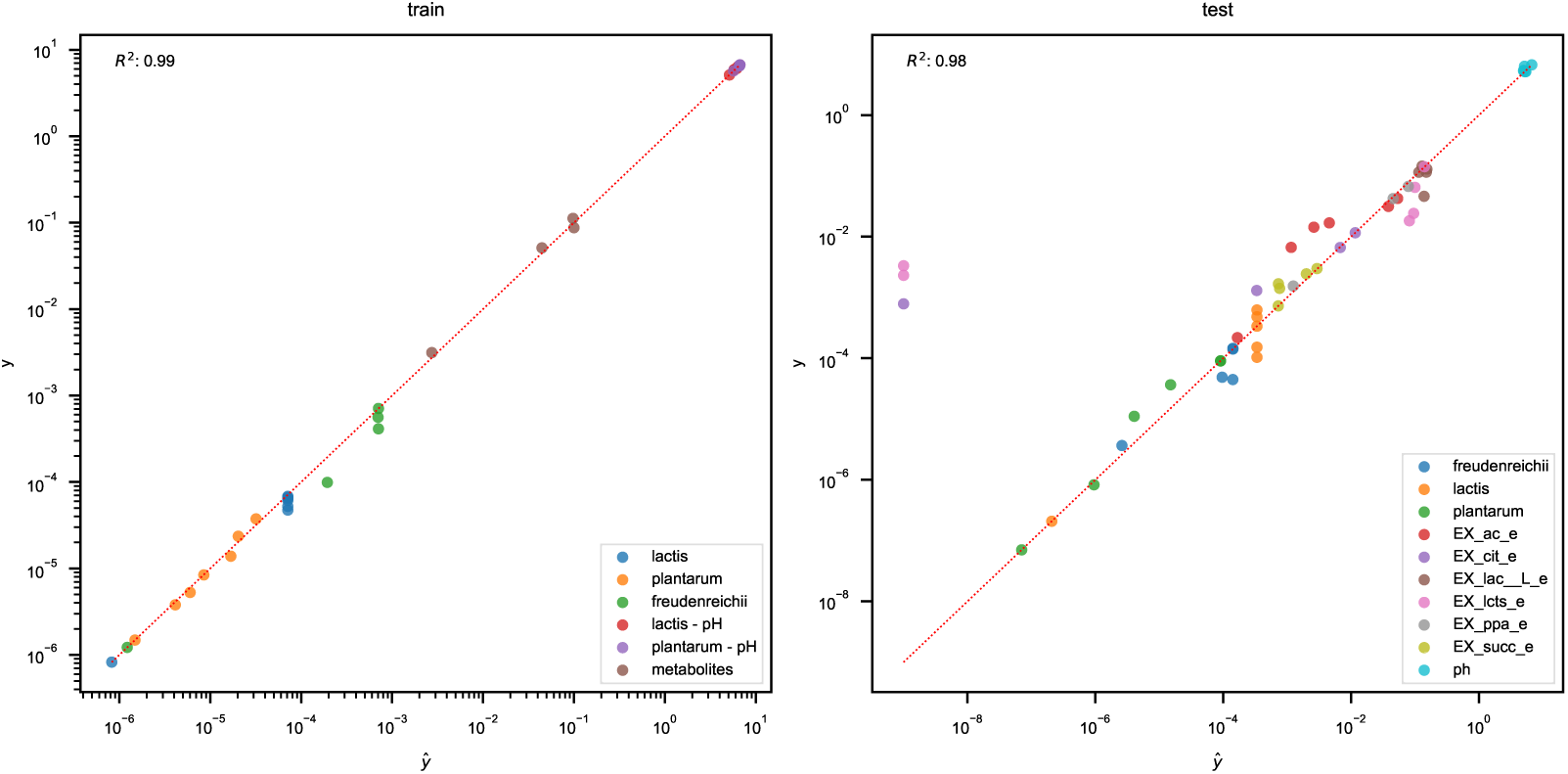
Model goodness of fit. All the model outputs (*y*^) are plotted versus their respective data value (*y*) in the training (i.e. the mono-culture data, left panel) and in the testing set (i.e. the co-culture data, right panel). Each point is colored according to the data type and the bissector is plotted (red dotted line). A global coefficient of determination is computed with the formula 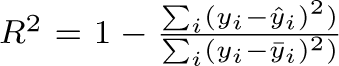 where *ȳ* is the average value of *y*. We can observe a very good agreement of the model in the training set, and a globally good agreement in the testing set, with 3 outliers on lactose and citrate.

**Figure S2:**
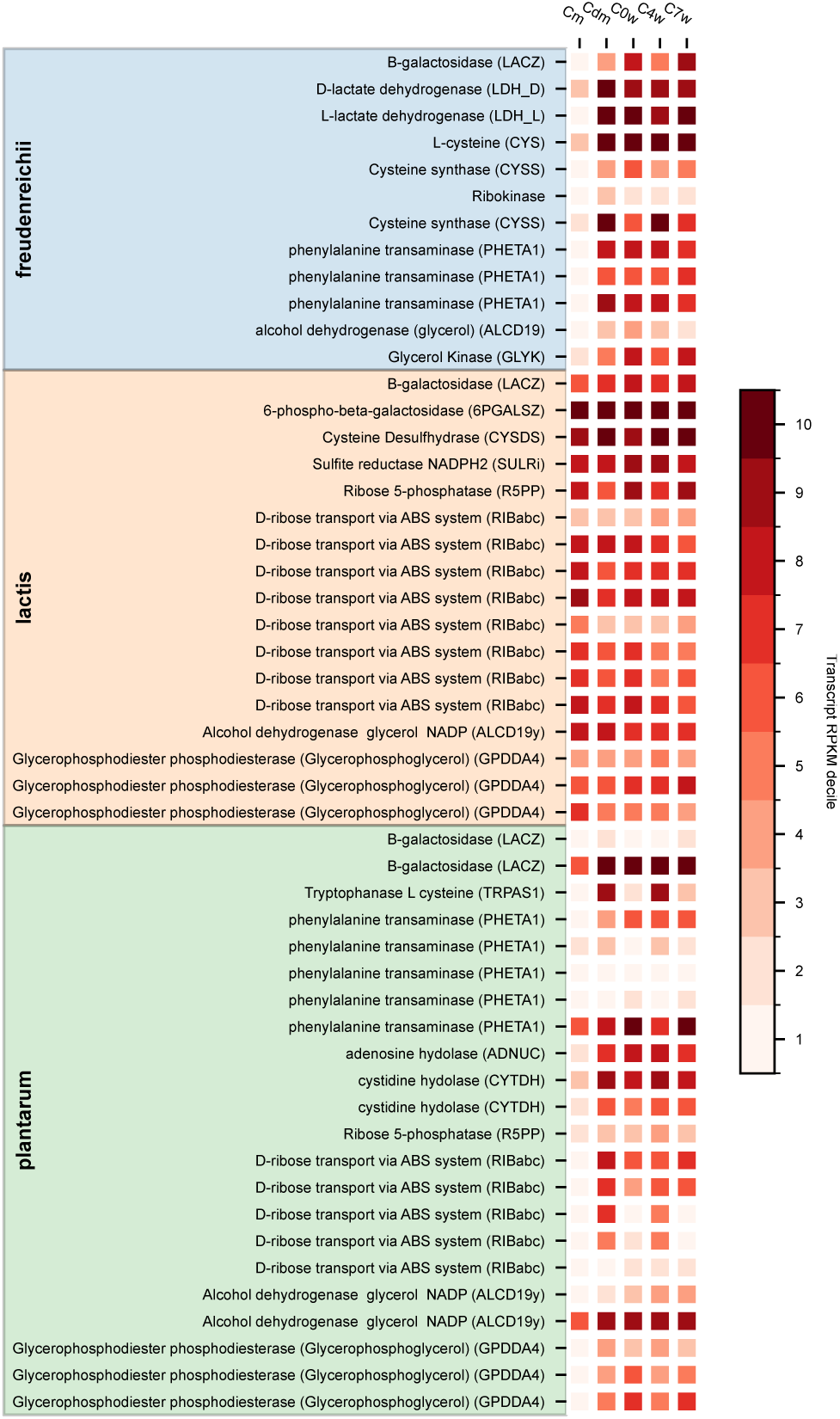
Gene expression with metatranscriptomic data. This heatmap displays the decile of the RPKM of gene transcript counts at a given time point (columns) and in a given micro-organism (the colored areas indicate to which bacteria the gene belongs). Then, the decile were color coded from highest deciles (dark red, highest expression in this micro-organism at this time point) to lowest decile (light red, low expression). The displayed genes were manually selected from the list of possible interactions raised by SMETANA: for a selected interaction metabolite, genes involved in production (in the donor), and in consumption (in the receiver) were retained for analysis. Selected interaction metabolites were *H*2*S*, ribose and phenylalanine. Additionally, genes involved in lactose consumption were added to this list. Abbreviations: Cm, molding (t = 19.5 hours); Cdm, demolding (t = 40 hours); C0w, start of ripening (t = 60 hours); C4w, Fourth week ripening (t = 732 hours); C7w, seventh week ripening (t = 1236 hours).

**Figure S3:**
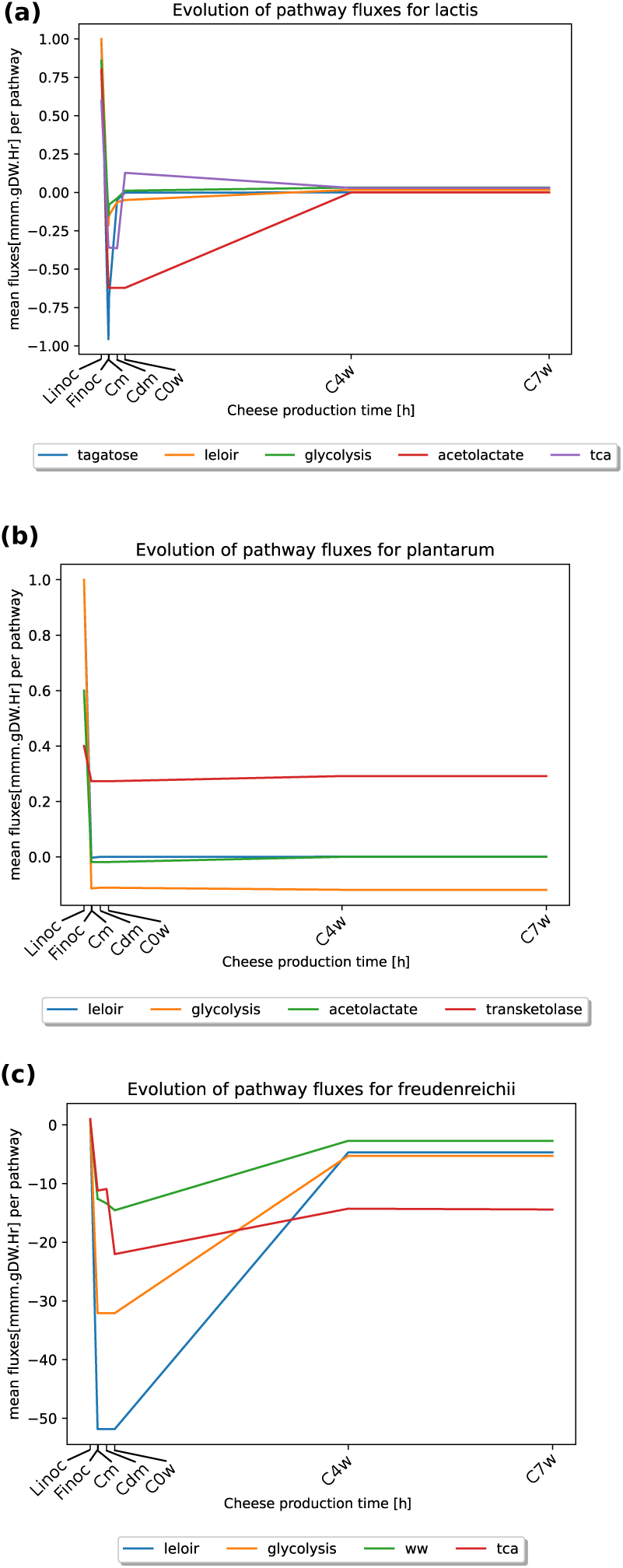
Metabolic pathway switches during cheese production. For each bacterium we ran a dFBA simulation and reaction fluxes per each pathways of interest in Figure 1 were retrieved at the inoculum of LAB (Linoc, t=0 hours), the inoculum of *P. freudenreichii* (Finoc, t=18 hours the molding (Cm, t=19.5 hours), demolding (Cdm, t=40hours), start of ripening (C0w, t=60 hours), fourth week ripening (C4w, t=732 hours) and seventh week ripening state (C7w, t=1236 hours). We normalized fluxes by the inoculum flux value and the average flux for each pathways is therefore represented.

## B Supplementary Material

### B.1 Pure culture data

#### B.1.1 Strain pure culture data

**Table S1:** Pure culture data used for the dynamic calibration of each strain

| GEM | Time | Bacterial density | pH |
| --- | --- | --- | --- |
| *P. freudenreichii* | 0 | 9.405e-07 | 6.76 |
|  | 8 | 1.221e-06 | 6.76 |
|  | 31 | 9.9e-05 | 6.55 |
|  | 48 | 0.000561 | 6.22 |
|  | 56 | 0.0004125 | 6.1 |
|  | 130 | 0.0007095 | 5.7 |
| *L. plantarum* | 0 | 1.485e-06 | 6.7 |
|  | 5 | 3.795e-06 | 6.68 |
|  | 7 | 5.28e-06 | 6.66 |
|  | 9 | 8.415e-06 | 6.645 |
|  | 14 | 1.386e-05 | 6.555 |
|  | 16 | 2.3595e-05 | 6.54 |
|  | 79 | 3.762e-05 | 6.705 |
| *L. lactis* | 0 | 8.25e-07 | 6.7 |
|  | 5 | 5.247e-05 | 6.49 |
|  | 7 | 6.831e-05 | 6.315 |
|  | 9 | 6.666e-05 | 6.075 |
|  | 14 | 6.468e-05 | 5.935 |
|  | 16 | 4.752e-05 | 8.865 |
|  | 79 | 6.171e-05 | 5.115 |

#### B.1.2 Acids dosage for *P. freudenreichii*

**Table S2:** Acids dosage data in g/L for *P. freudenreichii* .

| Compounds | Concentration $T_0$ | Concentration $T_f = 89$ h | Std |
| --- | --- | --- | --- |
| Lactate | 16.5 | 7.88 | 0.08 |
| Acetate | 0 | 3.07 | 0.02 |
| Succinate | 0 | 0.371 | 0.063 |
| Propionate | 0 | 88.31 | 0.02 |

### B.2 Manual refinement of GEMs

To ensure the production of extracellular compounds, we added manual modifications on intracellular bounds. These modifications are recapitulated in Table S3.

**Table S3:** Manual refinement of the GEMs and corresponding justifications.

| GEM | Reaction | Bound | Value | Justification |
| --- | --- | --- | --- | --- |
| *L. lactis* | ACTD2 | lower<br>upper | 0<br>0 | Allows the activation of the acetolactate pathway |
|  | ACLDC | lower<br>upper | 0<br>2 | Allows the activation of the acetolactate pathway |
|  | ACLS | lower<br>upper | 2<br>1000 | Allows the activation of the acetolactate pathway |
|  | LACZ | lower<br>upper | 0.0006<br>2 | Force the consumption of lactose |
| *L. plantarum* | PFL | lower<br>upper | 0<br>0 | Permit the production of both isomers of lactate from lactose. |
|  | LCTSt3ipp | lower<br>upper | 0<br>0 | Leads the flux of lactose into the Glycolysis pathway |
|  | ACTD2 | lower<br>upper | 0<br>0 | Allows the activation of the acetolactate pathway |
|  | GALUi | lower<br>upper | -30<br>10 | Permit the production of both isomers of lactate from lactose. |
|  | ACLS | lower<br>upper | 2<br>1000 | Allows the activation of the acetolactate pathway |
| *P. freudenreichii* | 6PGALSZ | lower<br>upper | 0<br>0 | Tagatose pathway was used |
|  | XYLI2 | lower<br>upper | 0<br>0 | Production of fructose from glucose instead of glucose-6-phosphate. Glycolysis was not activated |
|  | POX2 | lower<br>upper | 0<br>0 | Regulate flux of acetate |
|  | PPAKr | lower<br>upper | 0<br>0 | Inhibit acetate over production |
|  | LACZ | lower<br>upper | 0<br>5 | Allow consumption of lactose |
|  | PTAr | lower<br>upper | -1000<br>8 | Allow the appropriate ratio of acetate/propionate |
|  | 2131pyrpp | lower<br>upper | 8<br>1000 | Allow the appropriate ratio of acetate/propionate |

### B.3 Community experimental data

**Table S4:** Co-culture growth data used for testing community model predictions

| GEM | steps | Time (h) | Bacterial population ($\log_{10}$ CFU/g) |
| --- | --- | --- | --- |
| *P. freudenreichii* | Linoc | 0 | 6.1 |
|  | Finoc | 18 | 6.1 |
|  | Cm | 19.5 | 7.16 |
|  | C0w | 60 | 8.11 |
|  | C4w | 732 | 8.56 |
|  | C7w | 1236 | 8.59 |
| *L. plantarum* | Linoc | 0 | 5.2 |
|  | Finoc | 18 | 5.4 |
|  | Cm | 19.5 | 6.49 |
|  | C0w | 60 | 7.92 |
|  | C4w | 732 | 8.47 |
|  | C7w | 1236 | 8.47 |
| *L. lactis* | Linoc | 0 | 5.7 |
|  | Finoc | 18 | 7.5 |
|  | Cm | 19.5 | 8.78 |
|  | C0w | 60 | 9.06 |
|  | C4w | 732 | 8.95 |
|  | C7w | 1236 | 8.45 |

### B.4 Exchanged metabolites highlighted by computational approaches

#### B.4.1 MICOM

**Table S5:** Exchangeable metabolites highlighted by MICOM on the cheese bacterial community.

| Bigg ID | Metacyc ID | Name | Metacyc Ontology | Export fluxes | Import fluxes |
| --- | --- | --- | --- | --- | --- |
| acald_e | ACETALD | acetaldehyde | Aldehydes-Or-Ketones, Aldehydes | Pf; Ll | Lp |
| akg_e | 2-KETOGLUTARATE | 2-oxoglutarate | Acids, Organic-Acids | Pf | Lp |
| ala__D_e | D-ALANINE | D-ALANINE | All-Amino-Acids, Acids, Amino-Acids, Organic-Acids | Pf; Lp | Ll |
| ala__L_e | L-ALPHA-ALANINE | L-ALPHA-ALANINE | All-Amino-Acids, Acids, Amino-Acids, Organic-Acids | Pf; Lp | Ll |
| co2_e | CARBON-DIOXIDE | CARBON-DIOXIDE | Others, Others | Pf; Ll | Lp |
| coa_e | CO-A | co enzyme A | Groups, Others | Pf; Ll | Lp |
| fe2_e | FE+2 | Fe2+ | Ions, Inorganic-Ions, Cations | Lp | Pf; Ll |
| gly_e | GLY | Glycine | All-Amino-Acids, Acids, Amino-Acids, Organic-Acids | Lp | Pf; Ll |
| glyclt_e | GLYCOLATE | GLYCOLATE | Acids, Organic-Acids | Pf | Lp |
| gthox_e | OXIDIZED-GLUTATHIONE | glutathione disulfide | ORGANOSULFUR, All-Glutathiones | Lp | Ll |
| gthrd_e | GLUTATHIONE | glutathione | ORGANOSULFUR, All-Glutathiones, Thiols | Ll | Lp |
| h2o2_e | HYDROGEN-PEROXIDE | hydrogen peroxide | Peroxides, Others | Ll | Lp |
| h2o_e | WATER | H2O | Pseudo-Compounds, Others | Pf; Ll | Lp |
| h2s_e | HS | hydrogen sulfide | Ions, Anions, Inorganic-Ions | Ll | Lp; Pf |
| ile__L_e | ILE | isoleucine | All-Amino-Acids, Acids, Amino-Acids, Organic-Acids | Lp | Pf; Ll |
| lac__D_e | D-LACTATE | d-lactate | Acids, Organic-Acids | Pf | Ll; Lp |
| lac__L_e | L-LACTATE | l-lactate | Acids, Organic-Acids | Ll | Lp; Pf |
| mal__L_e | MAL | malate | Acids, Organic-Acids | Pf; Lp | Ll |
| nh4_e | AMMONIUM | ammonium | Ions, Inorganic-Ions, Cations | Lp | Pf; Ll |
| phe__L_e | PHE | phenylalanine | Aromatics, All-Amino-Acids, Acids, Amino-Acids, Organic-Acids, Organic-aromatic-compounds | Pf | Lp |
| pro__L_e | PRO | proline | All-Amino-Acids, Acids, Amino-Acids, Organic-Acids | Ll | Pf |
| ser__D_e | D-SERINE | serine | All-Amino-Acids, Acids, Amino-Acids, Organic-Acids | Ll | Lp; Pf |
| ser__L_e | SER | serine | All-Amino-Acids, Acids, Amino-Acids, Organic-Acids | Ll | Lp; Pf |
| succ_e | SUC | succinate | Acids, Organic-Acids | Ll | Pf |
| xan_e | XANTHINE | xanthine | Others, Others | Ll | Lp; Pf |

#### B.4.2 SMETANA

**Table S6:** Exchangeable metabolites highlighted by SMETANA on the cheese bacterial community.

| Bigg ID | Metacyc ID | Name | Metacyc Ontology | Export fluxes | Import fluxes |
| --- | --- | --- | --- | --- | --- |
| acald_e | ACETALD | acetaldéhyde | Aldehydes-Or-Ketones, Aldehydes | Pf<br>Pf<br>Lp<br>Ll | Ll<br>Lp<br>Ll<br>Lp |
| akg_e | 2-KETOGLUTARATE | 2-oxoglutarate | Acids, Organic-Acids | Lp<br>Pf | Pf<br>Lp |
| fum_e | FUM | fumarate | Acids, Organic-Acids | Pf | Lp |
| glyc_e | GLYCEROL | GLYCEROL | Alcohols, All-Carbohydrates, Sugar-alcohols, Carbohydrates | Lp<br>Ll<br>Ll | Pf<br>Pf<br>Lp |
| glyclt_e | GLYCOLATE | GLYCOLATE | Acids, Organic-Acids | Lp | Pf |
| h2s_e | HS | hydrogen sulfide | Ions, Anions, Inorganic-Ions | Lp<br>Pf<br>Ll | Pf<br>Lp<br>Lp |
| lac__D_e | D-LACTATE | d-lactate | Acids, Organic-Acids | Pf<br>Ll | Lp<br>Lp |
| lac__L_e | L-LACTATE | l-lactate | Acids, Organic-Acids | Ll | Lp |
| mal__L_e | MAL | malate | Acids, Organic-Acids | Pf<br>Lp<br>Pf<br>Ll | Ll<br>Ll<br>Lp<br>Lp |
| phe__L_e | PHE | phenylalanine | Aromatics, All-Amino-Acids, Acids, Amino-Acids<br>Organic-Acids, Organic-aromatic-compounds | Lp | Pf |
| rib__D_e | D-Ribopyranose | D-Ribopyranose | Aldehydes-Or-Ketones, All-Carbohydrates, Aldehydes, Carbohydrates | Ll<br>Lp<br>Pf<br>Ll | Pf<br>Pf<br>Lp<br>Lp |
| ser__D_e | D-SERINE | serine | All-Amino-Acids, Acids, Amino-Acids, Organic-Acids | Ll<br>Lp<br>Pf<br>Lp<br>Pf<br>Ll | Pf<br>Pf<br>Ll<br>Ll<br>Lp<br>Lp |
| succ_e | SUC | succinate | Acids, Organic-Acids | Ll<br>Lp | Pf<br>Pf |
| xan_e | XANTHINE | xanthine | Others, Others | Ll<br>Lp | Pf<br>Pf |

#### B.4.3 Metage2Metabo

**Table S7:**
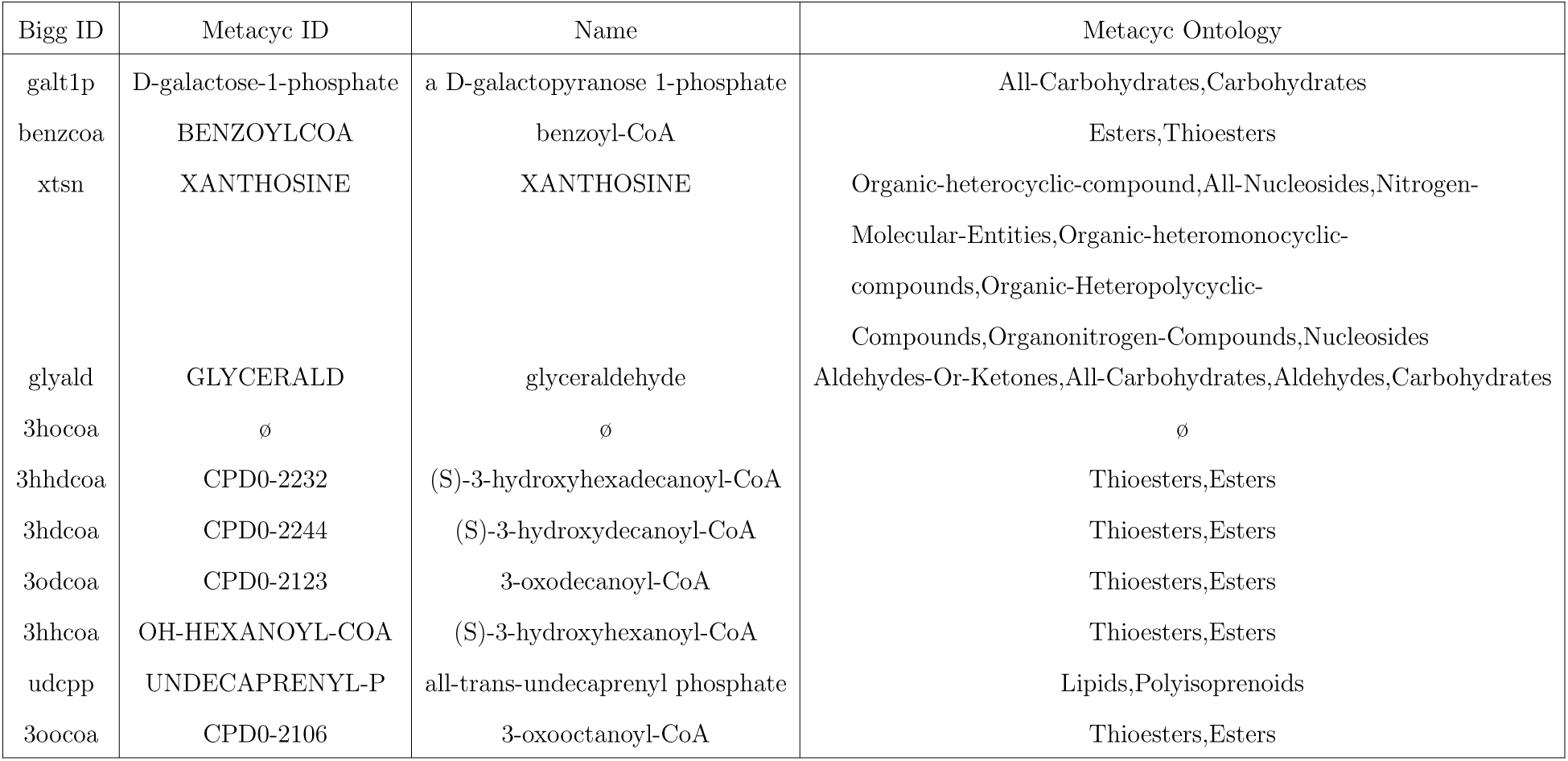
Metabolites predicted to be producible through metabolic cross-feeding only by Metage2Metabo

### B.5 Dynamic bounds on exchange reactions

In this section, we recapitulate and justify the regulation functions modulating the metabolite exchanges. For each GEM, the exchange reactions are listed, and their lower and upper bounds are specified.

#### L. lactis

- EX_lcts_e:

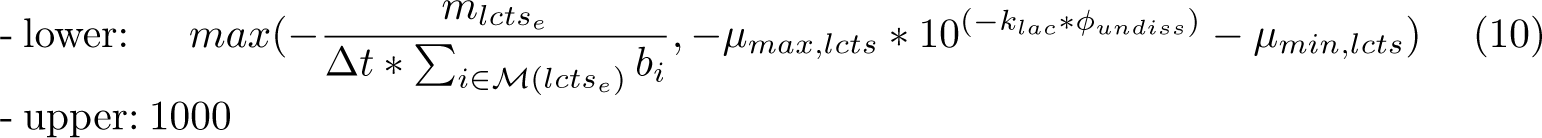 Justification: When lactose is depleted, the first term is activated, modeling uni-form sharing of available lactose among the set of bacteria *M*(*lac_L_*) metabolizing lactose. Otherwise, a regulation by the pH is added: the amount of undissociated lactic acid molecules is computed from lactate concentrations through the function *ϕ_undiss_* (see (7) for the expression of this function); then, an exponential increase from the value 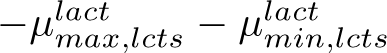 to the value 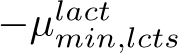 when the undissociated lactic acid concentration increases, with exponential rate *k_lac_*. The values of 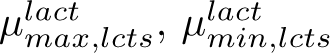 and *k_lac_* are inferred from co-culture experiments.
- EX_lac L_e, EX_lac D_e:

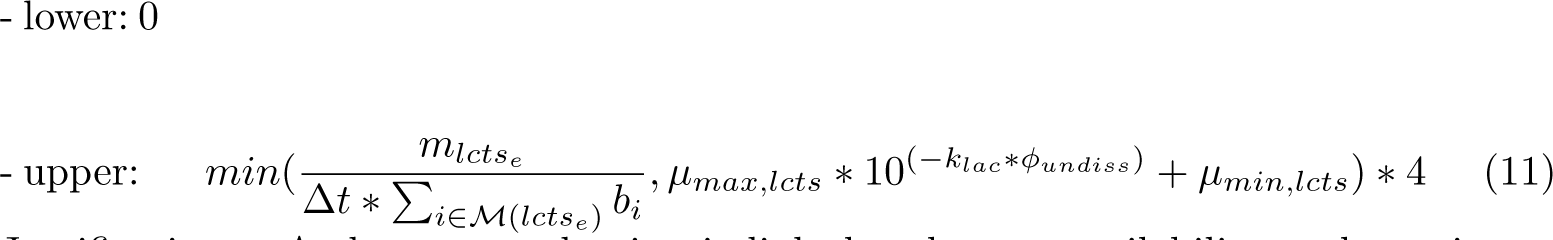 Justification: As lactate production is linked to lactose availability, a dynamic modulation for lactate export mimics lactose import (see eq. (12)), with a stoichiometric factor 4.0.
- EX_ac_e:

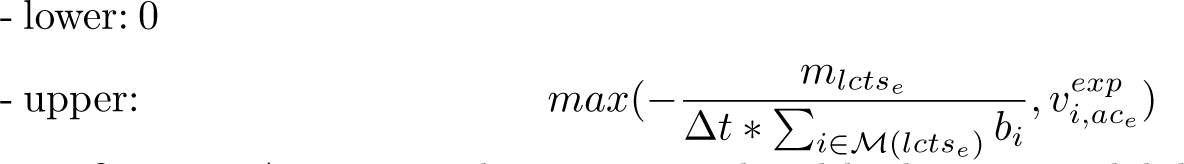 Justification: Acetate production is regulated by lactose availability when lactose is depleted, and is otherwise exported according to an intrinsic physiological export 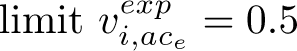.
- EX_diact_e, EX_btd_RR_e, EX_cit_e:

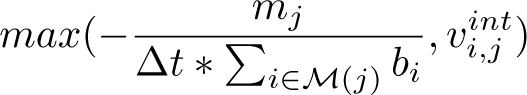 Justification: Usual consumption dynamic limitation (see eq. (6)). When substrate is not limited, import is constrained by intrinsic physiological limit modeled by *v^int^* = *−*8 for *j* =*L. lactis* and *j ∈ {EX*_*diact*_*e, EX*_*btd*_*RR*_*e}* and *v^int^* = *−*1 for *j* = *EX*_*cit*_*e*, and otherwise by nutrient availability. Available substrate is uniformly shared by consuming micro-organisms. Additionnaly, the lower bounds of *EX*_*glc D*_*e* (glucose), *EX*_*coa*_*e* (coenzyme A) and *EX*_*starch*1200_*e* (starch) were set to 0, since those compounds do not appear in milk composition.

#### L. plantarum

- EX_lcts_e:

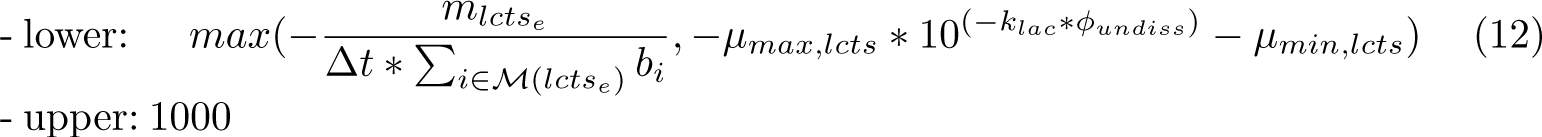 Justification: When lactose is depleted, the first term is activated, modeling uniform sharing of available lactose among the set of bacteria *M*(*lac_L_*) metabolizing lactose. Otherwise, a regulation by the pH is added: the amount of undissociated lactic acid molecules is computed from lactate concentrations through the function *ϕ_undiss_* (see (7) for the expression of this function); then, an exponential increase from the value 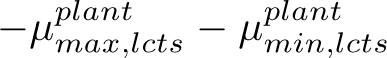 to the value 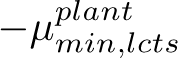 when the undissociated lactic acid concentration increases, with exponential rate *k_lac_*. The values of 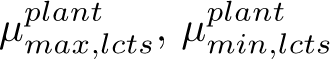 and *k_lac_* are inferred from co-culture experiments.
- EX_lac L_e, EX_lac D_e:

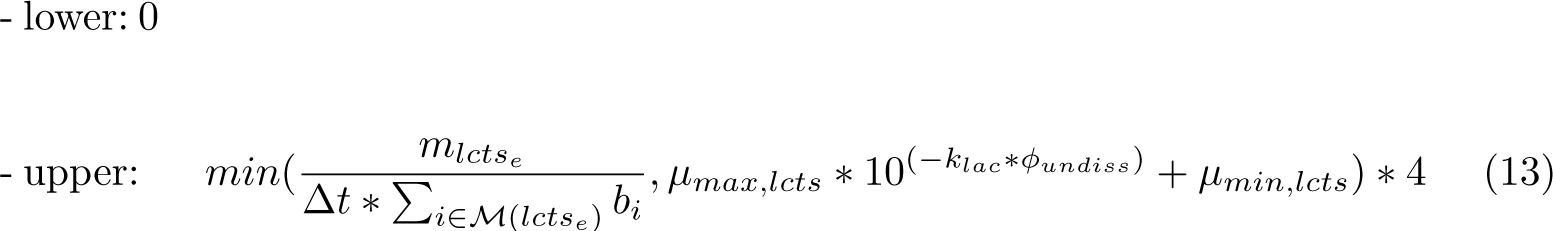 Justification: As lactate production is linked to lactose availability, a dynamic modulation for lactate export mimics lactose import (see eq. (12)), with a stoichiometric factor 4.0.
- EX_ac_e:

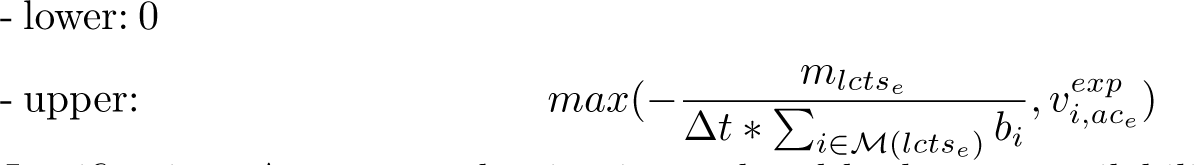 Justification: Acetate production is regulated by lactose availability when lactose is depleted, and is otherwise exported according to an intrinsic physiological export 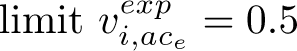.
- EX_btd_RR_e:

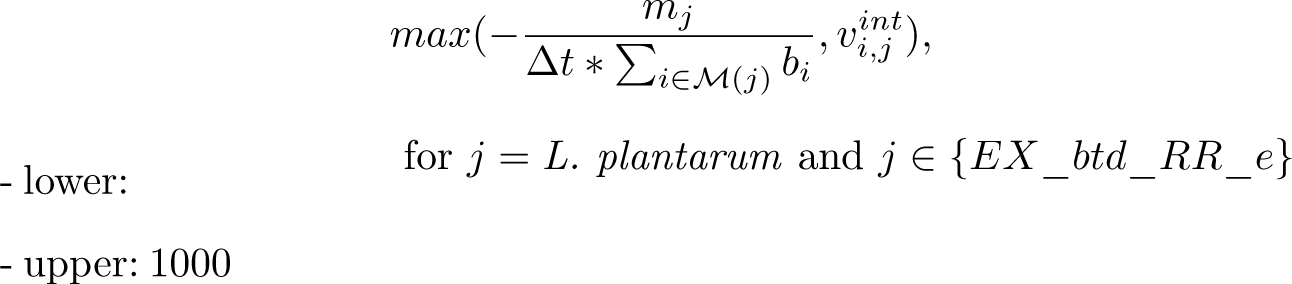 Justification: Usual consumption dynamic limitation (see eq. (6)). When substrate is not limited, import is constrained by intrinsic physiological limits modeled by 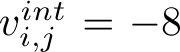 for *i* = *L. plantarum* and *j ∈ {EX*_*ac*_*e, EX*_*btd*_*RR*_*e}*, and 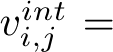 −1 for *j* = *EX*_*cit*_*e*, and otherwise by nutrient availability. Available substrate is uniformly shared by consuming micro-organisms. Additionally, the lower bounds of *EX*_*glc D*_*e* (glucose), *EX*_*gal*_*e* (galactose), *EX*_*coa*_*e* (coenzyme A) and *EX*_*dha*_*e* (dihydroxyacetone) were set to 0, since those compounds do not appear in milk composition.

#### P. freudenreichii

- EX_ser L_e: - lower: -12 - upper: 1000 Justification: Static consumption of serine
- EX_cit_e: - lower: -12 - upper: 1000 Justification: Static consumption of citrate
- EX_glu L_e: - lower: -12 - upper: 1000 Justification: Static consumption of glutamate
- EX_ala L_e: - lower: -12 - upper: 1000 Justification: Static consumption of alanine
- EX_lac L_e, EX_lac D_e:

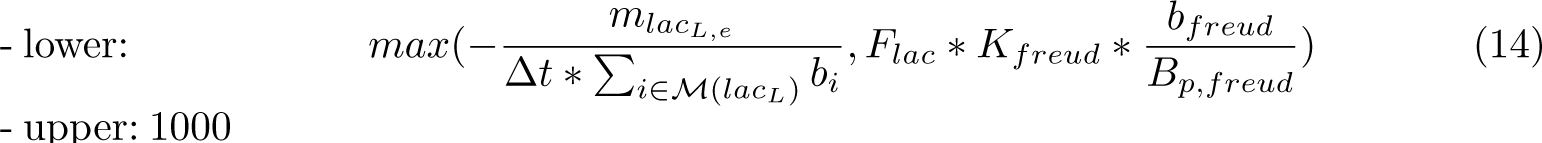 Justification: When lactate is depleted, the first term is activated, modeling uniform sharing of available lactate among the set of bacteria *M*(*lac_L_*) metabolizing L_lactate (resp. D_lactate). Otherwise, a flux *F_lac_*, estimated from metabolomic data in pure culture (see section B.7), is applied after modulation by the fraction 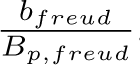, where *B_p,freud_* is the plateau biomass concentration value in pure culture *p,freud* and *b_freud_* is the current biomass density, and the factor *K_freud_*, which is inferred from the pure culture growth data. This modulation models a different order of magnitude of metabolism yield in pure or co-culture.
- EX_lcts_e:

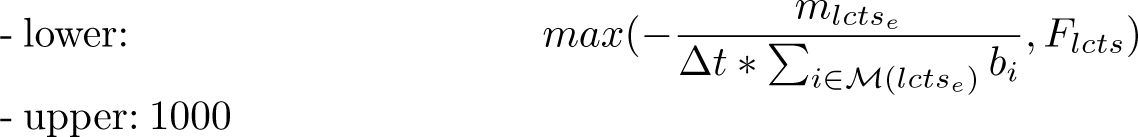 Justification: When lactose is depleted, the first term is activated, modeling uniform sharing of available lactose among the set of bacteria *M*(*lac_L_*) metabolizing lactose. Otherwise, a constant flux *F_lcts_*, estimated from metabolomic data in pure culture (see section B.7), is applied.
- EX_ac_e, EX_ppa_e,EX_succ_e:

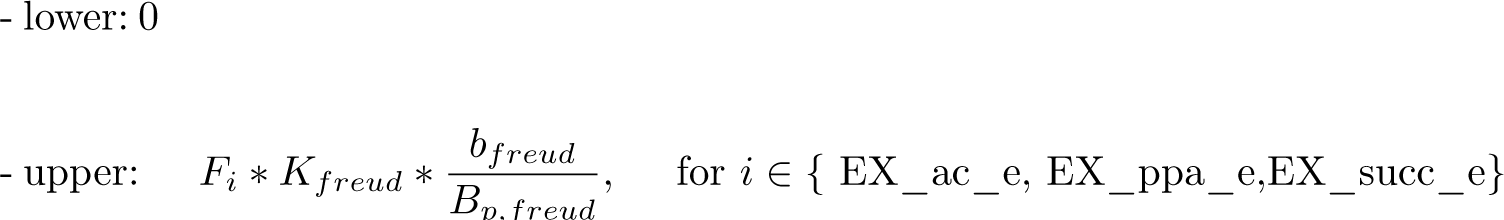 Justification: The maximal production rate is defined by *F_i_*, for *i ∈ {* EX_ac_e, EX_ppa_e,EX_succ_e*}*, which is estimated from metabolomic data in pure culture (see section B.7). The same modulation term than in 14 is applied, modeling a different order of magnitude of metabolism yield in pure or co-culture. Additionnaly, the lower bounds of *EX*_*starch*1200_*e* (starch) and *EX*_*dha*_*e* (dihydroxyacetone) were set to 0, since those compounds do not appear in milk composition.

### B.6 Dynamic bounds on intracellular reactions

When regulating metabolites involved in dynamic bounds for exchange reactions are depleted, the corresponding substrate import vanishes, inducing low metabolic fluxes across the whole GEM. These low metabolic fluxes can conflict with the fixed bounds on intracellular reactions introduced in Table S3, leading to infeasible FBA models. To avoid such infeasibility, we added additional dynamic bounds on the intracellular reactions. We stress that these dynamic bounds on intracellular reactions have a very different status than those on the exchange reactions: while the dynamic bounds on the exchange reactions drive the dynamics of the metabolic flux in the GEM according to substrate availability all along the simulation, the dynamics bounds on the intracellular reactions are only active when a substrate is depleted. They are “security” bounds that activate in case of vanishing substrate and weak metabolic activity, to avoid numerical infeasibility.

For each reaction contained in Table S3 indicating manually refined reactions, we add the following dynamic regulation

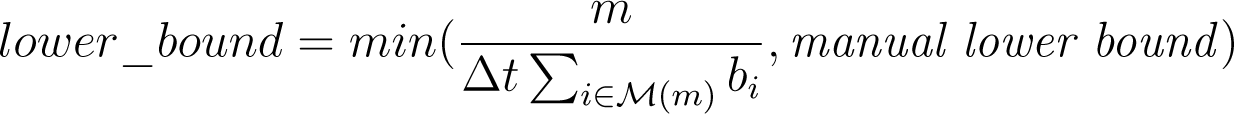

Hence, as the dynamic regulation is always positive, this regulation is active for strictly positive values of *manual lower bound*, the value indicated in Table S3. In that case, this regulation becomes active only when the metabolite *m* is depleted.

### B.7 Estimating bounds on metabolite production for *P. freudenreichii*

The dFBA equation in the pure culture experiment for *i* = *P. freudenreichii* reads,

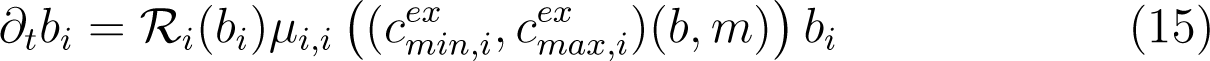

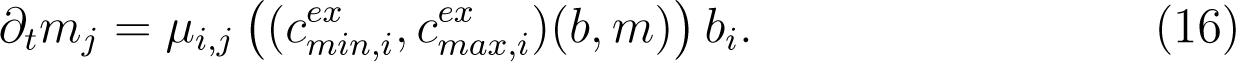

Assuming a constant production flux *µ_i,j_* and integrating (16) in time, we get

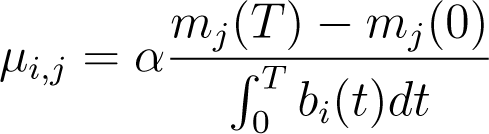

where *α* is a coefficient accounting for approximation errors.

We then evaluated *µ_i,j_* from initial and final metabolite concentrations, and numerical integration (trapeze scheme) of the biomass, taking *α* = 1.08. The resulting value is taken as lower bound for consumption and upper bound for production.

**Table S8:**
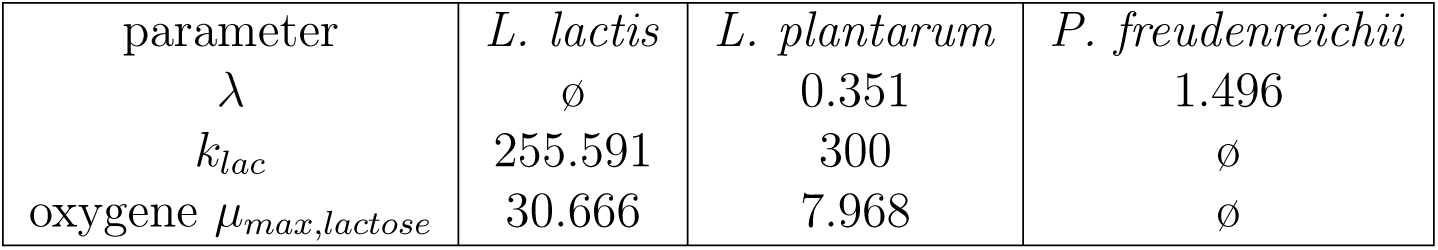
list of optimized parameters obtained after optimization and used in the fitted models.

### B.8 Parameter values

The optimized parameter values are recapitulated in Table S8. The remaining parameter values are recapitulated in Table S9.

**Table S9:**
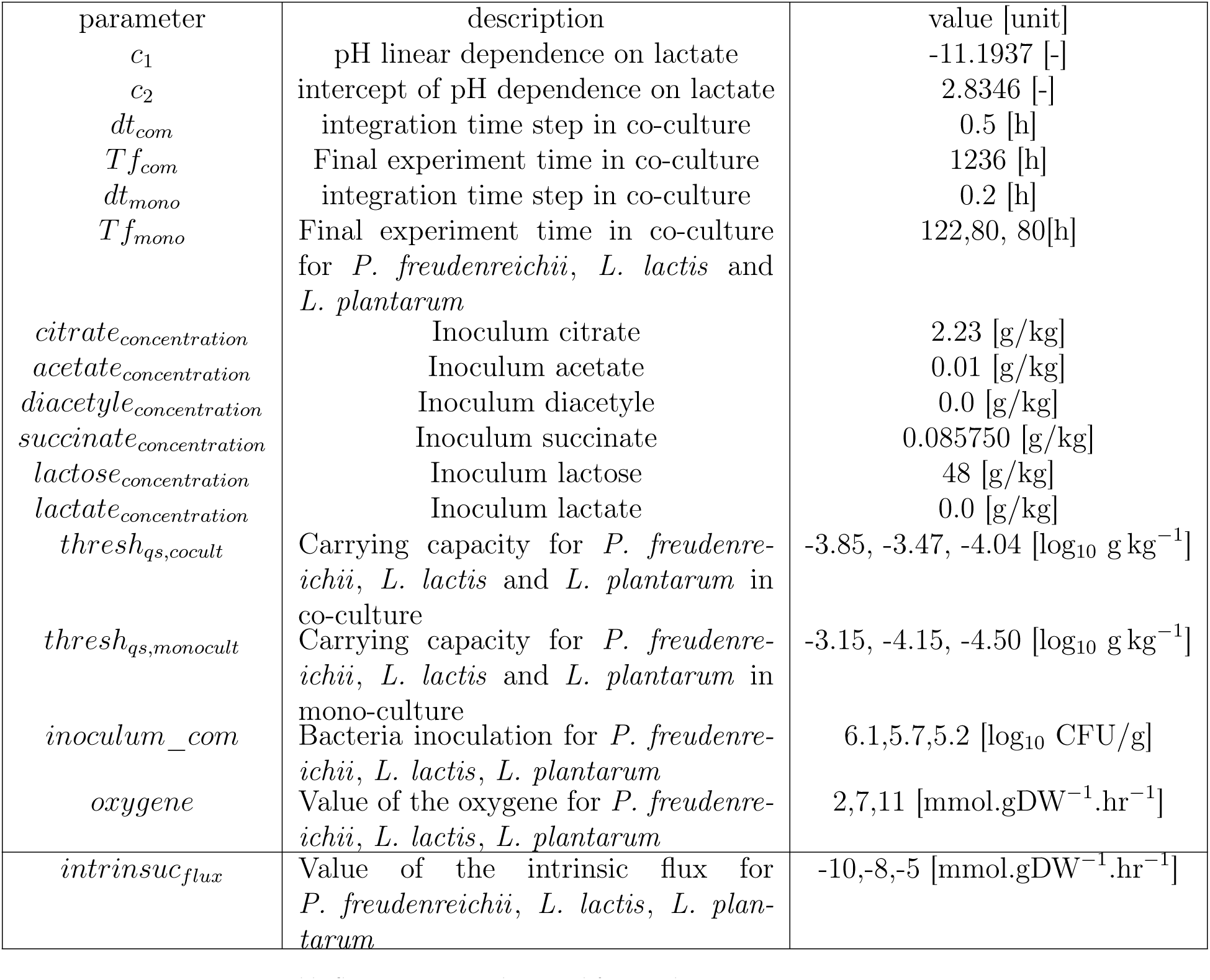
Parameter values used for simulations

## Footnotes

1 https://collection-cirmbia.fr/

2 https://escher.readthedocs.io/en/latest/escher-python.html

## References

1. Abedi, E. and Hashemi, S. M. B. (2020). Lactic acid production – producing microorganisms and substrates sources-state of art. Heliyon, 6(10):e04974.

2. Belcour, A., Frioux, C., Aite, M., Bretaudeau, A., Hildebrand, F., and Siegel, A. (2020). Metage2metabo, microbiota-scale metabolic complementarity for the identi1cation of key species. eLife.

3. Blasche, S., Kim, Y., Mars, R. A. T., Machado, D., Maansson, M., Kafkia, E., Milanese, A., Zeller, G., Teusink, B., Nielsen, J., Benes, V., Neves, R., Sauer, U., and Patil, K. R. (2021). Metabolic cooperation and spatiotemporal niche partitioning in a kefir microbial community. Nature Microbiology, 6(2):196–208.

4. Borghei, R., Mobasheri, M., and Sobati, T. (2021). Genome Scale Constraint-Based Metabolic Reconstruction of Propionibacterium Freudenreichii DSM 20271. Preprint from Research square, pages 1–29.

5. Cao, W., Aubert, J., Maillard, M.-B., Boissel, F., Leduc, A., Thomas, J.-L., Deutsch, S.-M., Camier, B., Kerjouh, A., Parayre, S., Harel-Oger, M., Garric, G., Thierry, A., and Falentin, H. (2021). Fine-Tuning of Process Parameters Modulates Specific Metabolic Bacterial Activities and Aroma Compound Production in Semi-Hard Cheese. Journal of Agricultural and Food Chemistry, 69(30):8511–8529.

6. Carroll, N. M., Ross, R. P., Kelly, S. M., Price, N. C., Sheehan, D., and Cogan, T. M. (1999). Characterization of recombinant acetolactate synthase from Leuconostoc lactis NCW1. Enzyme and Microbial Technology, 25(1-2):61–67.

7. Caspi, R., Billington, R., Fulcher, C. A., Keseler, I. M., Kothari, A., Krum-menacker, M., Latendresse, M., Midford, P. E., Ong, Q., Ong, W. K., Paley, S., Subhraveti, P., and Karp, P. D. (2018). The MetaCyc database of metabolic pathways and enzymes. Nucleic Acids Research, 46(D1):D633– D639.

8. Crow, V. L. (1986). Metabolism of aspartate by propionibacterium freudenreichii subsp. shermanii: Effect on lactate fermentation. Applied and Environmental Microbiology, 52(2):359–365.

9. Dank, A., van Mastrigt, O., Boeren, S., Lillevang, S. K., Abee, T., and Smid, E. J. (2021). Propionibacterium freudenreichii thrives in microaerobic conditions by complete oxidation of lactate to CO2. Environmental Microbiology, 23(6):3116–3129.

10. Deborde, C. and Boyaval, P. (2000). Interactions between pyruvate and lactate metabolism in Propionibacterium freudenreichii subsp. shermanii: In vivo 13C nuclear magnetic resonance studies. Applied and Environmental Microbiology, 66(5):2012–2020.

11. Diener, C., Gibbons, S. M., and Resendis-Antonio, O. (2020a). MICOM: Metagenome-Scale Modeling To Infer Metabolic Interactions in the Gut Microbiota. mSystems, 5(1).

12. Diener, C., Gibbons, S. M., and Resendis-Antonio, O. (2020b). MICOM: Metagenome-Scale Modeling To Infer Metabolic Interactions in the Gut Microbiota. mSystems, 5(1).

13. Ebrahim, A., Lerman, J. A., Palsson, B. O., and Hyduke, D. R. (2013). COBRApy: COnstraints-Based Reconstruction and Analysis for Python. BMC Systems Biology, 7(1):74.

14. Falentin, H., Deutsch, S.-M., Jan, G., Loux, V., Thierry, A., Parayre, S., Maillard, M.-B., Dherbécourt, J., Cousin, F. J., Jardin, J., Siguier, P., Couloux, A., Barbe, V., Vacherie, B., Wincker, P., Gibrat, J.-F., Gaillardin, C., and Lortal, S. (2010). The Complete Genome of Propionibacterium freudenreichii CIRM-BIA1T, a Hardy Actinobacterium with Food and Probiotic Applications. PLoS ONE, 5(7):e11748.

15. Fang, X., Lloyd, C. J., and Palsson, B. O. (2020). Reconstructing organisms in silico: genome-scale models and their emerging applications. Nature Reviews Microbiology, pages 1–13.

16. Galimberti, A., Bruno, A., Agostinetto, G., Casiraghi, M., Guzzetti, L., and Labra, M. (2021). Fermented food products in the era of globalization: tradition meets biotechnology innovations. Current Opinion in Biotechnology, 70:36–41.

17. Gomez, J. A., Höffner, K., and Barton, P. I. (2014). Dfbalab: a fast and reliable matlab code for dynamic flux balance analysis. BMC bioinformatics, 15(1):1–10.

18. Harris, C. R., Millman, K. J., van der Walt, S. J., Gommers, R., Virtanen, P., Cournapeau, D., Wieser, E., Taylor, J., Berg, S., Smith, N. J., Kern, R., Picus, M., Hoyer, S., van Kerkwijk, M. H., Brett, M., Haldane, A., del Río, J. F., Wiebe, M., Peterson, P., Gérard-Marchant, P., Sheppard, K., Reddy, T., Weckesser, W., Abbasi, H., Gohlke, C., and Oliphant, T. E. (2020). Array programming with NumPy. Nature, 585(7825):357–362.

19. Hunter, J. D. (2007). Matplotlib: A 2D Graphics Environment. Computing in Science & Engineering, 9(3):90–95.

20. King, Z. A., Dräger, A., Ebrahim, A., Sonnenschein, N., Lewis, N. E., and Palsson, B. O. (2015). Escher: A Web Application for Building, Sharing, and Embedding Data-Rich Visualizations of Biological Pathways. PLoS Computational Biology.

21. Kleerebezem, M., Boekhorst, J., Van Kranenburg, R., Molenaar, D., Kuipers, O. P., Leer, R., Tarchini, R., Peters, S. A., Sandbrink, H. M., Fiers, M. W., Stiekema, W., Klein Lankhorst, R. M., Bron, P. A., Hoffer, S. M., Nierop Groot, M. N., Kerkhoven, R., De Vries, M., Ursing, B., De Vos, W. M., and Siezen, R. J. (2003). Complete genome sequence of Lactobacillus plantarum WCFS1. Proceedings of the National Academy of Sciences of the United States of America, 100(4):1990–1995.

22. Langmead, B., Trapnell, C., Pop, M., and Salzberg, S. (2009). Langmead b, trapnell c, pop m, salzberg sl.. ultrafast and memory-efficient alignment of short dna sequences to the human genome. genome biol 10: R25. Genome biology, 10:R25.

23. Lieven, C., Beber, M. E., Olivier, B. G., Bergmann, F. T., Ataman, M., Babaei, P., Bartell, J. A., Blank, L. M., Chauhan, S., Correia, K., Diener, C., Dräger, A., Ebert, B. E., Edirisinghe, J. N., Faria, J. P., Feist, A. M., Fengos, G., Fleming, R. M. T., García-Jiménez, B., Hatzimanikatis, V., Helvoirt, W. v., Henry, C. S., Hermjakob, H., Herrgård, M. J., Kaafarani, A., Kim, H. U., King, Z., Klamt, S., Klipp, E., Koehorst, J. J., König, M., Lakshmanan, M., Lee, D.-Y., Lee, S. Y., Lee, S., Lewis, N. E., Liu, F., Ma, H., Machado, D., Mahadevan, R., Maia, P., Mardinoglu, A., Medlock, G. L., Monk, J. M., Nielsen, J., Nielsen, L. K., Nogales, J., Nookaew, I., Palsson, B. O., Papin, J. A., Patil, K. R., Poolman, M., Price, N. D., Resendis-Antonio, O., Richelle, A., Rocha, I., Sánchez, B. J., Schaap, P. J., Sheriff, R. S. M., Shoaie, S., Sonnenschein, N., Teusink, B., Vilaça, P., Vik, J. O., Wodke, J. A. H., Xavier, J. C., Yuan, Q., Zakhartsev, M., and Zhang, C. (2020). MEMOTE for standardized genome-scale metabolic model testing. Nature Biotechnology, 38(3):272–276.

24. Loux, V., Mariadassou, M., Almeida, S., Chiapello, H., Hammani, A., Buratti, J., Gendrault, A., Barbe, V., Aury, J. M., Deutsch, S. M., Parayre, S., Madec, M. N., Chuat, V., Jan, G., Peterlongo, P., Azevedo, V., Le Loir, Y., and Falentin, H. (2015). Mutations and genomic islands can explain the strain dependency of sugar utilization in 21 strains of Propionibacterium freudenreichii. BMC Genomics, 16(1):1–19.

25. Machado, D., Andrejev, S., Tramontano, M., and Patil, K. R. (2018). Fast automated reconstruction of genome-scale metabolic models for microbial species and communities. bioRxiv, page 223198.

26. Mahadevan, R., Edwards, J. S., and Doyle, F. J. (2002). Dynamic Flux Balance Analysis of Diauxic Growth in Escherichia coli. Biophysical Journal, 83(3):1331–1340.

27. Makhlouf, A. (2006). Méthodologie pour l’optimisation dynamique multicritère d’un procédé discontinu alimenté: application à la production bactérienne d’arômes laitiers. PhD thesis, Institut National Polytechnique de Lorraine.

28. Mannaa, M., Han, G., Seo, Y.-S., and Park, I. (2021). Evolution of Food Fermentation Processes and the Use of Multi-Omics in Deciphering the Roles of the Microbiota. Foods, 10(11):2861.

29. McKinney, W. et al. (2010). Data structures for statistical computing in python. In Proceedings of the 9th Python in Science Conference, volume 445, pages 51–56. Austin, TX.

30. Miyoshi, A., Rochat, T., and Gratadoux, J.-j. (2003). Miyoshi. Genetics and Molecular Research, 2(4):348–359.

31. Noronha, A., Modamio, J., Jarosz, Y., Guerard, E., Sompairac, N., Preciat, G., Daníelsdóttir, A. D., Krecke, M., Merten, D., Haraldsdóttir, H. S., Heinken, A., Heirendt, L., Magnúsdóttir, S., Ravcheev, D. A., Sahoo, S., Gawron, P., Friscioni, L., Garcia, B., Prendergast, M., Puente, A., Rodrigues, M., Roy, A., Rouquaya, M., Wiltgen, L., Žagare, A., John, E., Krueger, M., Kuperstein, I., Zinovyev, A., Schneider, R., Fleming, R. M. T., and Thiele, I. (2018). The Virtual Metabolic Human database: integrating human and gut microbiome metabolism with nutrition and disease. Nucleic Acids Research, 47(D1):D614–D624.

32. Orth, J. D., Thiele, I., and Palsson, B. O. (2010). What is flux balance analysis? Nature Biotechnology, 28(3):245–248.

33. Palles, T., Beresford, T., Condon, S., and Cogan, T. M. (1998). Citrate metabolism in Lactobacillus casei and Lactobacillus plantarum. Journal of Applied Microbiology, 85(1):147–154.

34. Palsson, B. O. and Varma, A. (1994). Metabolic Flux Balancing: Basic Concepts, Scientific and Practical Use. Biotechnology, 12(October):994– 998.

35. Putri, G. H., Anders, S., Pyl, P. T., Pimanda, J. E., and Zanini, F. (2022). Analysing high-throughput sequencing data in Python with HTSeq 2.0. Bioinformatics, 38(10):2943–2945.

36. Quatravaux, S., Remize, F., Bryckaert, E., Colavizza, D., and Guzzo, J. (2006). Examination of Lactobacillus plantarum lactate metabolism side effects in relation to the modulation of aeration parameters. Journal of Applied Microbiology, 101(4):903–912.

37. R Core Team (2021). R: A Language and Environment for Statistical Computing. R Foundation for Statistical Computing, Vienna, Austria.

38. Rau, M. H. and Zeidan, A. A. (2018). Constraint-based modeling in microbial food biotechnology. Biochemical Society Transactions, 46(2):249–260.

39. Robinson, M. D., McCarthy, D. J., and Smyth, G. K. (2009). edgeR: a Bioconductor package for differential expression analysis of digital gene expression data. Bioinformatics, 26(1):139–140.

40. Roissart, H. d. (DL 1994). Bactéries lactiques : aspects fondamentaux et technologiques / H. de Roissart, F. M. Luquet, coordonnateurs. Lorica, Uriage.

41. Römer, M., Eichner, J., Dräger, A., Wrzodek, C., Wrzodek, F., and Zell, A. (2016). ZBIT bioinformatics toolbox: A web-platform for systems biology and expression data analysis. PLoS ONE.

42. Smid, E. J. and Kleerebezem, M. (2014). Production of aroma compounds in lactic fermentations. Annual Review of Food Science and Technology, 5(1):313–326.

43. Somerville, V., Grigaitis, P., Battjes, J., Moro, F., and Teusink, B. (2021). Use and limitations of genome-scale metabolic models in food microbiology. Current Opinion in Food Science.

44. Swindell, S. R., Benson, K. H., Griffin, H. G., Renault, P., Ehrlich, S. D., and Gasson, M. J. (1996). Genetic manipulation of the pathway for diacetyl metabolism in Lactococcus lactis. Applied and Environmental Microbiology, 62(7):2641–2643.

45. Tamang, J. P., Shin, D.-H., Jung, S.-J., and Chae, S.-W. (2016a). Functional Properties of Microorganisms in Fermented Foods. Frontiers in Microbiology, 7:578.

46. Tamang, J. P., Watanabe, K., and Holzapfel, W. H. (2016b). Review: Diversity of microorganisms in global fermented foods and beverages. Frontiers in Microbiology, 7(MAR).

47. Thierry, A., Deutsch, S. M., Falentin, H., Dalmasso, M., Cousin, F. J., and Jan, G. (2011). New insights into physiology and metabolism of Propionibacterium freudenreichii. International Journal of Food Microbiology, 149(1):19–27.

48. Turgay, M., Bachmann, H.-P., Irmler, S., von Ah, U., Frö Hlich- Wyder, M.-T., Falentin, H., Deutsch, S.-M., Jan, G., and Thierry, A. (2020). Propionibacterium spp. and Acidipropionibacterium spp. In <u>Reference Module in Food Science</u>, Reference Module in Food Science. Elsevier.

49. Vallenet, D., Calteau, A., Dubois, M., Amours, P., Bazin, A., Beuvin, M., Burlot, L., Bussell, X., Fouteau, S., Gautreau, G., Lajus, A., Langlois, J., Planel, R., Roche, D., Rollin, J., Rouy, Z., Sabatet, V., and Médigue, C. (2019). MicroScope: an integrated platform for the annotation and exploration of microbial gene functions through genomic, pangenomic and metabolic comparative analysis. Nucleic Acids Research.

50. Van Rooijen, R. J., Van Schalkwijk, S., and De Vos, W. M. (1991). Molecular cloning, characterization, and nucleotide sequence of the tagatose 6phosphate pathway gene cluster of the lactose operon of Lactococcus lactis. Journal of Biological Chemistry, 266(11):7176–7181.

51. Van Rossum, G. and Drake, F. L. (2009). Python 3 Reference Manual. CreateSpace, Scotts Valley, CA.

52. Wes McKinney (2010). Data Structures for Statistical Computing in Python. In Stéfan van der Walt and Jarrod Millman, editors, Proceedings of the 9th Python in Science Conference, pages 56 – 61.

53. Wick, R. R., Judd, L. M., Gorrie, C. L., and Holt, K. E. (2017a). Completing bacterial genome assemblies with multiplex MinION sequencing. Microbial Genomics, 3(10):1–7.

54. Wick, R. R., Judd, L. M., Gorrie, C. L., and Holt, K. E. (2017b). Unicycler: Resolving bacterial genome assemblies from short and long sequencing reads. PLoS Computational Biology, 13(6):1–22.

55. Widyastuti, Y. R., and Febrisiantosa, A. (2014). The Role of Lactic Acid Bacteria in Milk Fermentation. Food and Nutrition Sciences, 05(04):435– 442.

56. Zelezniak, A., Andrejev, S., Ponomarova, O., Mende, D. R., Bork, P., and Patil, K. R. (2015). Metabolic dependencies drive species co-occurrence in diverse microbial communities. Proceedings of the National Academy of Sciences, 112(20):6449–6454.

57. Özcan, E., Seven, M., Şirin, B., Çakır, T., Nikerel, E., Teusink, B., and Öner, E. T. (2020). Dynamic co-culture metabolic models reveal the fermentation dynamics, metabolic capacities and interplays of cheese starter cultures. Biotechnology and Bioengineering, 118(1):223–237.

